# APOE is a presynaptic protein that accumulates with age and modulates neurotransmitter release

**DOI:** 10.64898/2026.04.20.719736

**Authors:** Sarayut Phasuk, Kyla B. Tooley, Julianna L. Sun, Vishwajeeth Pagala, Gustavo Palacios, Sean Deats, Gaven Garland, Laura L. Robinson, Xusheng Wang, Bonn Belingon, Jenn Cook, Haiyan Tan, Ankhbayar Lkhagva, Zuo-Fei Yuan, Long Wu, Amanda Johnson, Mazdak Bradberry, Camenzind G. Robinson, Anthony A. High, Ron Korstanje, Jason D. Vevea

## Abstract

The synaptic vesicle (SV) cycle is the fastest membrane trafficking and protein sorting process in biology. It underlies neuronal communication and cognition, yet synaptic function declines during normal aging, increasing vulnerability to neurologic disease. How the SV cycle is maintained across the lifespan of a complex organism remains unclear. Here, we used wild-type mice (C57BL/6J) to define the age- and sex-stratified molecular landscape of SVs and identified apolipoprotein E (APOE) as an abundant presynaptic protein further enriched in aged female samples. Super-resolution imaging, cell-type selective expression, and protease protection assays demonstrate that APOE originates from astroglia and associates with the cytosolic face of SVs. Using iGluSnFR and pHluorin optophysiology, we find that both decreased and increased APOE levels impair neurotransmission during stimulus trains. Together, these findings place APOE at the synapse and establish it as a cell-nonautonomous regulator of the SV cycle.

## Introduction

The synaptic vesicle (SV) cycle includes organelle biogenesis **(Bolz & Haucke, 2025)**, docking and fusion **(Kaeser & Regehr, 2017)**, endocytosis **(Imoto & Watanabe, 2025)**, and recycling **(Ivanova & Cousin, 2022)**. Across milliseconds to seconds, this cycle orchestrates rapid lipid and protein sorting to drive neurotransmitter release while completely dismantling and reforming the SV. Synaptic transmission therefore depends on robust organelle homeostasis mechanisms that act rapidly and consistently throughout life. Synaptic dyshomeostasis can appear physiologically as a loss of function at individual synapses, anatomically as a loss of synapse number, and behaviorally as cognitive impairment **(Brito et al., 2023)**. Consistent with this framework, previous studies observe impaired synaptic function **(Deupree, Bradley, & Turner, 1993; M. Wang et al., 2011)**, and reduced synapse number in multiple brain regions with advanced age in naturally aged rodents and non-human primates **(Morrison & Baxter, 2012)**. Unlike age-related cognitive decline, neurodegenerative diseases like Alzheimer’s disease (AD) drive accelerated cognitive impairment. Soluble toxic protein oligomers disrupt synaptic function early, with synapse loss emerging as disease progresses but preceding bulk neuronal death **(Davies, Mann, Sumpter, & Yates, 1987; Shankar et al., 2008)**. Across reports, synapse loss is the best pathological correlate of cognitive impairment **(DeKosky & Scheff, 1990; Scheff, Price, Schmitt, & Mufson, 2006; Terry et al., 1991)**. Despite extensive evidence for age- and disease-related synaptic dysfunction and loss, we lack a molecular investigation documenting key synaptic machinery like the SV across the lifespan of organisms.

Synaptosomes are well established nerve terminal preparations from brain tissue that retain presynaptic and postsynaptic components and have long served as a valuable platform for studies of age-related synaptic biology. Using rodent models, these studies routinely report age-associated reductions of multiple pre- and postsynaptic proteins relative to younger controls **(Canas, Duarte, Rodrigues, Kofalvi, & Cunha, 2009; Remesal et al., 2025; VanGuilder, Yan, Farley, Sonntag, & Freeman, 2010; Vegh et al., 2014)**. Yet synaptosomes are compositionally heterogeneous and contain mitochondria, endoplasmic reticulum (ER), and the plasma membrane (PM), in addition to pre- and postsynaptic components. This heterogeneity limits interpretation because age-related synaptic decline may reflect defects in specific compartments, including SVs, that cannot be resolved in mixed preparations. For example, the ‘poisoned t-SNARE’ (or ‘SNAP accumulation’) hypothesis proposes that impaired presynaptic sorting during SV recycling allows t-SNAREs to persist on newly formed SVs, rendering SNARE machinery less competent for future rounds of fusion **(Truckenbrodt et al., 2018)**. Something a mixed preparation that includes the PM cannot resolve. Just as proteins must be rapidly sorted to regenerate SVs, lipids are also tightly regulated and play essential roles in the SV cycle. For example, cholesterol comprises approximately 40 mol% of the SV lipidome. **(Breckenridge, Morgan, Zanetta, & Vincendon, 1973)**. SV cholesterol helps organize membrane proteins like the synaptophysin-synaptobrevin 2 (SYP-SYB2) interaction **(Mitter et al., 2003)**, or it supports protein function as in the case of the SV V-ATPase **(Yoshinaka, Kumanogoh, Nakamura, & Maekawa, 2004)**. Supporting a causal link between cholesterol homeostasis and synaptic performance, cognitive impairment is a core symptom of Niemann-Pick Disease Type C (NPC), and NPC1 knockout (KO) neurons show SV recycling defects that can be improved by adding back cholesterol **(Wasser, Ertunc, Liu, & Kavalali, 2007)**.

Testing such compartment-specific models requires biochemically defined sub-synaptic fractions, but these preparations are technically demanding and time-consuming, which has historically limited throughput. Nevertheless, foundational molecular characterization of the SV and post-synaptic density (PSD) proteomes were established in the mid-2000s **(Peng et al., 2004; Takamori et al., 2006)** and later refined using modern mass spectrometry techniques **(Distler et al., 2014; Taoufiq et al., 2020)**. However, direct SV-focused aging datasets were absent until recently. Gao et al. examined WT mouse synaptosomes and crude SV preparations using ultracentrifugation to find eight SV proteins moderately upregulated at 10 months of age **(Gao et al., 2025)**. These studies all have strengths but share important limitations. Most reports sample only two or three ages across the rodent lifespan, rely on heterogeneous synaptosomes preparations, or focus on animals that are not yet at advanced age. In addition, studies of synaptic function with age either do not mention sex as a biological variable or only examine one sex. However, sex may shape synaptic density and its decline with age. Rat cortex shows age-by-sex differences in spine density, and human cortex shows sex differences in synaptic density **(Alonso-Nanclares, Gonzalez-Soriano, Rodriguez, & DeFelipe, 2008; Markham & Juraska, 2002)**. The SV2A PET findings however, remain disputed with one study reporting no sex effect with age, while another finds region-specific declines in synaptic density **(Michiels et al., 2021; Toyonaga et al., 2024)**. Nevertheless, outside of age, female sex is among the strongest risk factors for dementia, making it a critical variable to consider **(Fang et al., 2025; Niccoli & Partridge, 2012)**.

To address these gaps, we performed a sex-stratified, age-dependent analysis of SV composition in wild-type (WT) C57BL/6J mice. Our goal was to define how this rapidly recycled organelle is maintained during aging and to determine whether age-dependent shifts in SV composition could explain synaptic vulnerability with age. We hypothesized that aging alters SV proteome and lipidome homeostasis by driving selective accumulation or depletion of key vesicle components. We combined high-plex TMT proteomics with untargeted and targeted lipidomics on highly enriched SVs isolated at 3, 6, 12, 18, 24, and 30 months from male and female mice (n = 4 per sex per age). This lifespan- and sex-stratified scale was only enabled by the recent validation of a rapid SV immunoprecipitation workflow (SV-RIP) **(Bradberry et al., 2022; Bradberry, Peters-Clarke, Shishkova, Chapman, & Coon, 2023)**. This design defines age- and sex-dependent changes in SV composition and also provides a direct test of compartment-specific models, including the “poisoned t-SNARE” hypothesis.

After establishing a reproducible SV-RIP multiomics workflow, we performed life-course profiling to test for age-dependent changes in SV composition. Our approach detected >3300 proteins and >350 lipid species finding remarkable stability of the SV proteome and lipidome across the mouse lifespan. Hierarchical clustering uncovered a set of proteins that increase with age, with larger increases in females. This pattern matches the two strongest risk factors for dementia: advanced age and female sex. This cluster contained proteins linked to neurodegeneration in humans, including the key complement initiator C1q, and lipid trafficking protein apolipoprotein E (APOE). We prioritized APOE because its isoforms modulate age-related cognitive function **(Shinohara et al., 2016)**, it is the strongest genetic determinant for late-onset AD **(Liu, Liu, Kanekiyo, Xu, & Bu, 2013)**, and its risk effects are stronger in females than in males **(Altmann, Tian, Henderson, Greicius, & Alzheimer’s Disease Neuroimaging Initiative, 2014)**.

We identify mouse APOE as an abundant presynaptic protein in intact brain tissue throughout the hippocampus and cortex. Biochemical and fluorescence assays show that APOE associates with SVs and adopts a cytosol-facing orientation on the vesicle surface. In neuron–astroglia co-cultures with astroglia-restricted expression, astroglia-derived APOE accumulates at presynaptic terminals. Using optophysiology in WT, APOE KO, and astroglial APOE overexpression (OE) conditions, we find that both APOE loss and APOE excess reduce glutamate release probability and impair sustained neurotransmission during stimulus trains. Together, these results establish APOE as a cell-nonautonomous age- and sex-regulated presynaptic protein that modulates the SV cycle.

## Results

### Preparation and validation of synaptic vesicle rapid immunoprecipitation (SV-RIP) workflow

We isolated SVs based largely on an SV-RIP workflow as previously described **(Bradberry et al., 2022)**. SV-RIP relies on a monoclonal antibody to the conserved synaptic vesicle glycoprotein 2 (SV2) protein **(Buckley & Kelly, 1985)**. We varied brain homogenate (BH) clarification speeds (low/middle/high) and buffer tonicity (iso- vs hypotonic). SV-RIP remained robust across conditions, so we selected a middle speed with hypotonic buffer for subsequent experiments (**Supplemental Figure 1A-D**). Incubating clarified BH with SV2-coupled magnetic beads yielded a highly enriched SV fraction (**Figure 1A**). Total protein staining and targeted immunoblots revealed highly enriched SV proteins with low background and depletion of post-synaptic protein 95 (PSD95) and voltage-dependent anion channel (VDAC), markers of the post-synapse and mitochondria, respectively (**Figure 1B-C**). Ficin protease-based elution of bound material from the beads revealed small vesicles with an outer diameter averaging 55 nm by negative stain electron microscopy (EM), consistent with SV size (**Figure 1D-E**). Confident that SV-RIP isolates and enriches SVs from BH, we used data dependent acquisition and a standard spectral counting approach to compare protein eluted from IgG control beads and SV2 coated magnetic beads. We found clear enrichment of canonical SV proteins including v-ATPase subunits, synaptotagmin 1 (SYT1), synaptophysin (SYP), and SV2 isoforms. The IgG control beads bound cytoskeletal and RNA binding elements (**Figure 1F**). We then used our untargeted lipidomics pipeline and recovered lipids reflecting known abundant SV lipids, particularly enriched for phosphatidylcholine (PC) species and docosahexaenoic acid (DHA)-containing lipids (**Figure 1G**) **(Takamori et al., 2006)**. Together, these orthogonal validations confirm that SV-RIP cleanly and robustly isolates intact SVs from BH and is compatible with quantitative proteomic and lipidomic profiling, enabling the lifespan-scale analyses described next.

**Figure 1.**
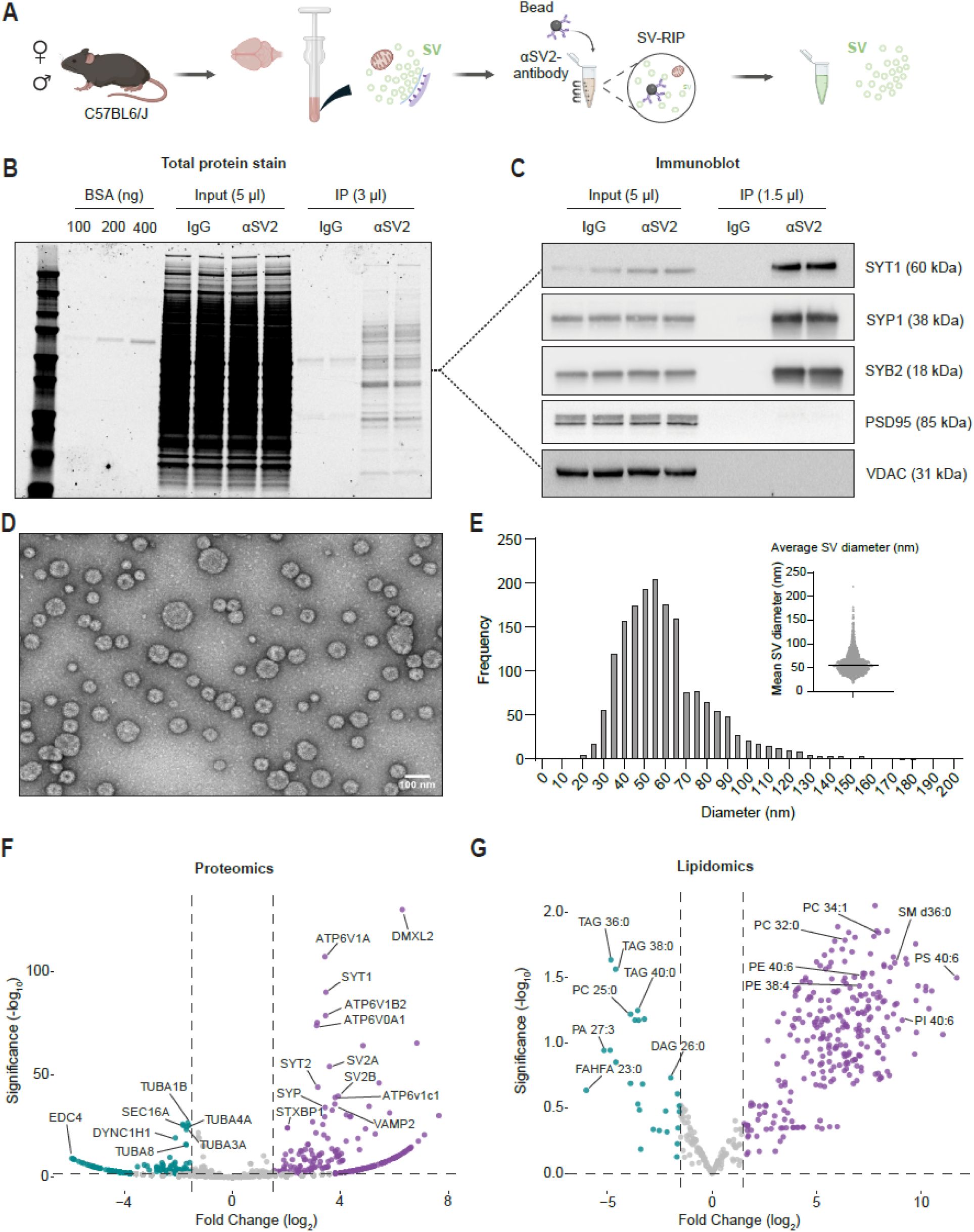
Preparation and validation of synaptic vesicle rapid isolation protocol (SV-RIP). **A.** Schematic workflow of synaptic vesicle rapid immunoprecipitation (SV-RIP). The whole brain of C57BL6/J mice was Dounce homogenized and Input collected. Homogenate was incubated with αSV2- or IgG-conjugated magnetic beads and collected with a magnetic stand after several wash steps. **B.** Total protein was stained with GelCode Blue for BSA (100, 200, 400 ng), IgG Input, SV2 Input, IgG-IP, and SV2-IP. **C.** Immunoblot analysis of IgG Input, SV2 Input, IgG-IP, and SV2-IP for synaptic vesicle markers (SYT1, SYP1, SYB2) and other organelle markers (PSD95, VDAC) to assess the purity of RIP-isolated synaptic vesicles. **D.** Transmission electron microscopy (TEM) of isolated synaptic vesicles. **E.** Distribution of synaptic vesicle diameter measured from TEM using SAMJ **(Garcia-Lopez-de-Haro et al., 2025)**, demonstrating an average diameter of ∼50 nm. **F-G.** Proteomics and lipidomics comparing fold enrichment between IgG and SV2 coated beads, analyzed using spectral counts and VolcaNoseR web app **(Goedhart & Luijsterburg, 2020)**.

### Multiplexed mouse synaptic vesicle proteomics reveals broad compositional stability across lifespan in both sexes

We validated our 18-plex TMT proteomics pipeline using young (3 month) male and female mice. Proteomics and immunoblots confirmed enrichment of SV proteins and depletion of other compartments (e.g., post-synapse, lysosome, Golgi, mitochondria, lipids droplet) (**Supplemental Figure 2A-B**). We then performed SV-RIP on male and female C57BL/6J mice, from 3, 6, 12, 18, 24, and 30 months of age (n = 4 per sex per age) (**Figure 2A**). All mice were alert, mobile, and healthy in appearance, with no signs of kyphosis or malnutrition, consistent with normal aging and showed no evidence of overt chronic age-related disease. In both sexes, body weight increased from 3 to 12 months and then plateaued from 12 to 30 months (**Supplemental Figure 2C**). Non-fasting blood glucose levels did not differ between young (3-month) and old (30-month) mice, indicative of healthy aged animals (**Supplemental Figure 2D**). One fraction of the isolated SV preparation was analyzed by 18-plex TMT-based proteomics using the in-house JUMP software analysis pipeline, and another fraction was used for follow-up immunoblotting **(Li et al., 2021; X. Wang et al., 2014)** (**Figure 2B; Supplemental Figure 2E**). Our sensitive TMT proteomics pipeline detected 3339 proteins from every sample analyzed, shown ranked by their abundance (**Figure 2C**). All samples were well correlated and broadly represented proteins related to the SV cycle using SynGO (**Supplemental Figure 3A-B**) **(Koopmans et al., 2019)**. Analyzing the top 1500 most abundant proteins (representing >90% of detected signal) by PCA plot revealed no clear grouping by sex or age (**Figure 2D**). We conclude the SV proteome is broadly stable throughout the mouse lifespan. We graphed by rank abundance curated lists of abundant SV proteins (**Figure 2E**), v-ATPase subunits (**Figure 2F**), and SV transporters (**Figure 2G**), finding no significant changes between young and old mice of either sex. We next sought to investigate SV protein levels in BH and validate our proteomics results using targeted immunoblots. Probing the BH for global levels of SYT1, SYP, and SYB2, we detected no changes across age or sex (**Figure 2H-I**). Analyzing the SV fraction by immunoblot revealed no changes in SYP or SYT1, but a significant decrease in SYB2 in males at 24 and 30 months of age (**Figure 2J-K**). These results support the proteomics dataset and further indicate a modest but significant decrease in SYB2 by immunoblot. This effect is likely below the detection threshold of our global proteomics analysis following multiple-testing correction and false discovery rate (FDR) control. Overall, our results demonstrate, for the first time, that the SV proteome is remarkably stable throughout the mouse lifespan.

**Figure 2.**
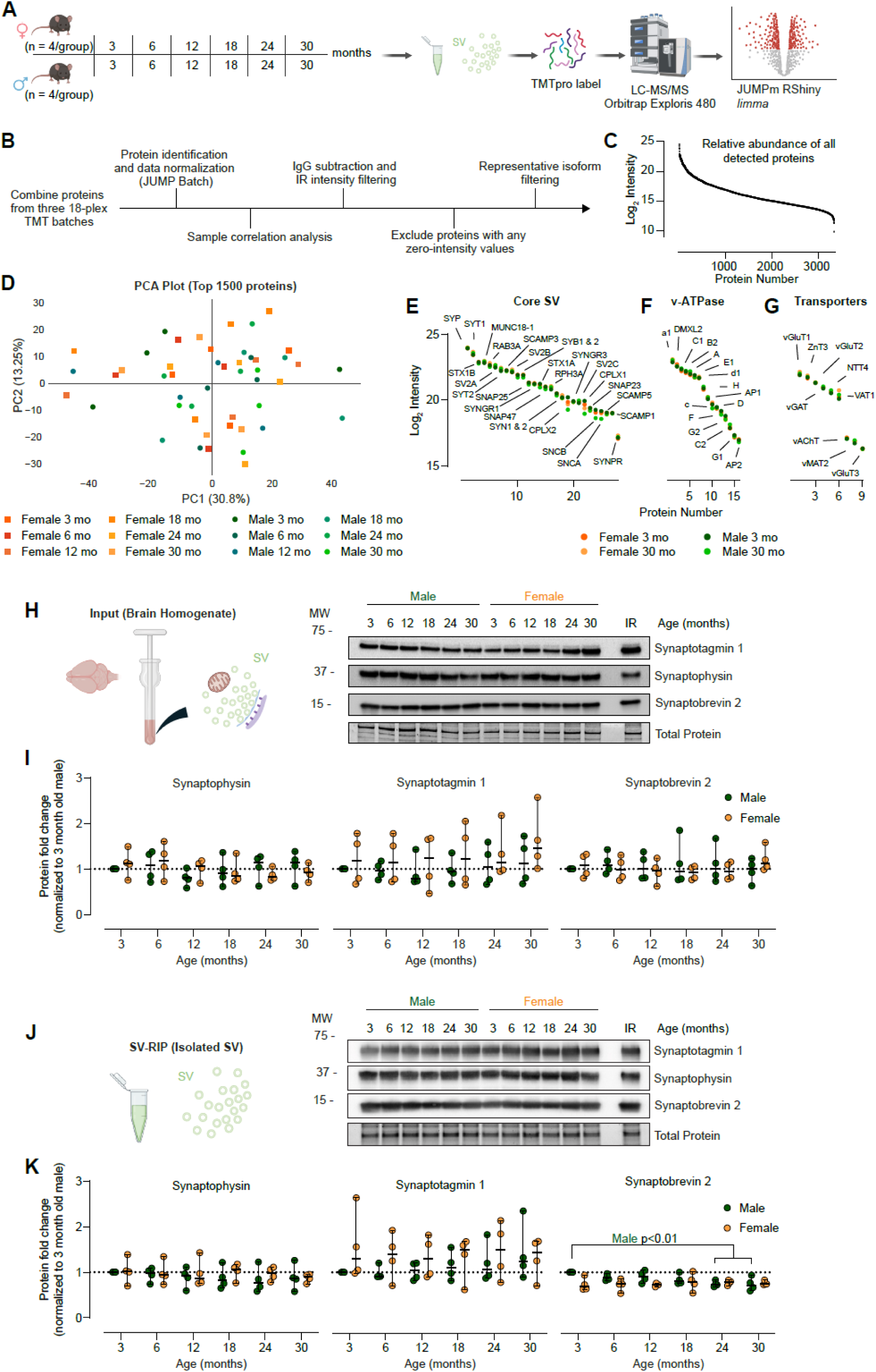
Multiplexed mouse synaptic vesicle proteomics reveals broad compositional stability across lifespan in both sexes. **A.** Experimental workflow for proteomic analysis of SVs. SVs were isolated from the whole brain of male and female C57BL6/J mice at 3-, 6-, 12-, 18-, 24-, and 30-months of age by SV-RIP and divided into 4 aliquots (n=4/group, N=48). One aliquot was taken for TMTpro Multiplexed Proteomics. **B.** Proteomics data analysis pipeline. Proteins from all batches were combined, with identification and data normalization performed with JUMP Batch. Sample correlation analysis was performed to identify potential outliers, followed by IgG subtraction and IR intensity filtering. Any proteins with one or more zero-intensity values were excluded from further analysis. Representative isoform filtering was performed, leaving 3339 proteins for downstream analyses. **C.** Rank ordered log2 intensity of all 3339 proteins detected by TMT Proteomics. **D.** Principal component analysis for all samples demonstrates no distinct clustering of by age or sex for the top 1500 most abundant proteins. **E.** Rank ordered log2 intensity of Core SV proteins from 3- and 30-month-old males and females show consistent protein levels with age for both sexes. **F.** Rank ordered log2 intensity of V-ATPase proteins from 3- and 30-month-old males and females show consistent protein levels with age for both sexes. **G.** Rank ordered log2 intensity of transporter proteins from 3- and 30-month-old males and females show consistent protein levels with age for both sexes. **H.** Representative immunoblot of 3-, 6-, 12-, 18-, 24-, and 30-month male and female Input samples and an internal reference (IR) for synaptic proteins SYT1, SYP, SYB2, and Total Protein. **I.** Immunoblot quantitation of 3-, 6-, 12-, 18-, 24-, and 30-month male and female Input samples, normalized to 3-month-old males. Data are displayed as median ± SEM and analyzed by 2-way ANOVA with Tukey’s multiple comparisons test (p>0.05). **J.** Representative immunoblot of 3-, 6-, 12-, 18-, 24-, and 30-month male and female Isolated SV samples and an internal reference (IR) for synaptic proteins SYT1, SYP, SYB2, and Total Protein. **K.** Immunoblot quantitation of 3-, 6-, 12-, 18-, 24-, and 30-month male and female Isolated SV samples, normalized to 3-month-old males. Data are displayed as median ± SEM and analyzed by 2-way ANOVA with Dunnett’s multiple comparisons test for SYB2 and with Tukey’s comparisons test for SYP and SYT1 (** p≤0.01).

### Apolipoprotein E is an abundant synaptic vesicle protein and accumulates with age in females

We performed limma differential analysis leveraging our in-house JUMPshiny application **(Ritchie et al., 2015; Zhang et al., 2025)**. Age is a gradual process, and our multi-timepoint design allowed us to model protein abundance as a function of age across the mouse lifespan. After applying FDR correction, 30 proteins showed a significant linear association with age. We then performed hierarchical clustering via k-means using the complexHeatmap R package for these 30 proteins and found they resolved best into 4 clusters (**Figure 3A**) **(Gu, Eils, & Schlesner, 2016)**. Cluster 1 includes proteins whose association with SVs increases with age, independent of sex. Cluster 2 includes proteins that increase with age but do so more strongly in females than in males. Cluster 3 contains a single protein that decreases with age in female SVs. Cluster 4 includes proteins whose association with SVs decreases with age, independent of sex (**Figure 3B**). Among these 30 significantly altered proteins, a striking feature is that many are non-neuronal proteins but with established roles in synaptic maintenance and homeostasis. Notably, Cluster 2 is enriched for proteins linked to age-related disease, including AD (e.g., C1q, APOE) **(Corder et al., 1993; Hong et al., 2016)**. These patterns support our working hypothesis that specific factors accumulate on, or are depleted from, SVs with age, and that these effects are amplified in females consistent with the higher incidence of dementia in older women compared with men **(Fang et al., 2025; Niccoli & Partridge, 2012)**. We next ranked these 30 proteins by their relative abundance and performed STRING pathway enrichment analysis for associated biological processes **(Szklarczyk et al., 2025)** (**Figure 3C**). APOE was the most abundant protein in this set and exhibited the highest connectivity in STRING, with most interactions occurring among proteins within the same cluster. Hierarchical clustering in Figure 3A was driven solely by abundance profiles, whereas STRING leverages curated protein–protein interaction networks; strikingly, despite these orthogonal inputs, both approaches converged on a similar Cluster 2 module, increasing confidence in the robustness and biological relevance of this finding. We next performed post hoc pairwise comparisons between 3- and 30-month SV proteomes in males and females to highlight sex-specific aging differences and complement our across-age analysis using limma (**Figure 3D**). We then carried out Gene Ontology (GO) enrichment analysis for biological processes (**Supplemental Figure 4A**) **(Yu, Wang, Han, & He, 2012)**. Complement activation emerged as the top enriched process. Finally, we performed follow up immunoblotting to quantify APOE and C1q in BH (input) and SV fraction. In both sexes, APOE levels did not increase globally in BH; instead, APOE showed an age-dependent increase in SV association, with a larger accumulation on SVs from females, supporting a specific vesicle interaction. For C1q, males and females showed an increase in protein levels within BH, along with increased C1q in the SV fraction (**Figure 3E-H**). These results indicate that APOE is an abundant presynaptic protein (**Supplemental Figure 4B**) whose increased association with SVs reflects a specific, age-dependent interaction rather than a global age-related increase in APOE abundance.

**Figure 3.**
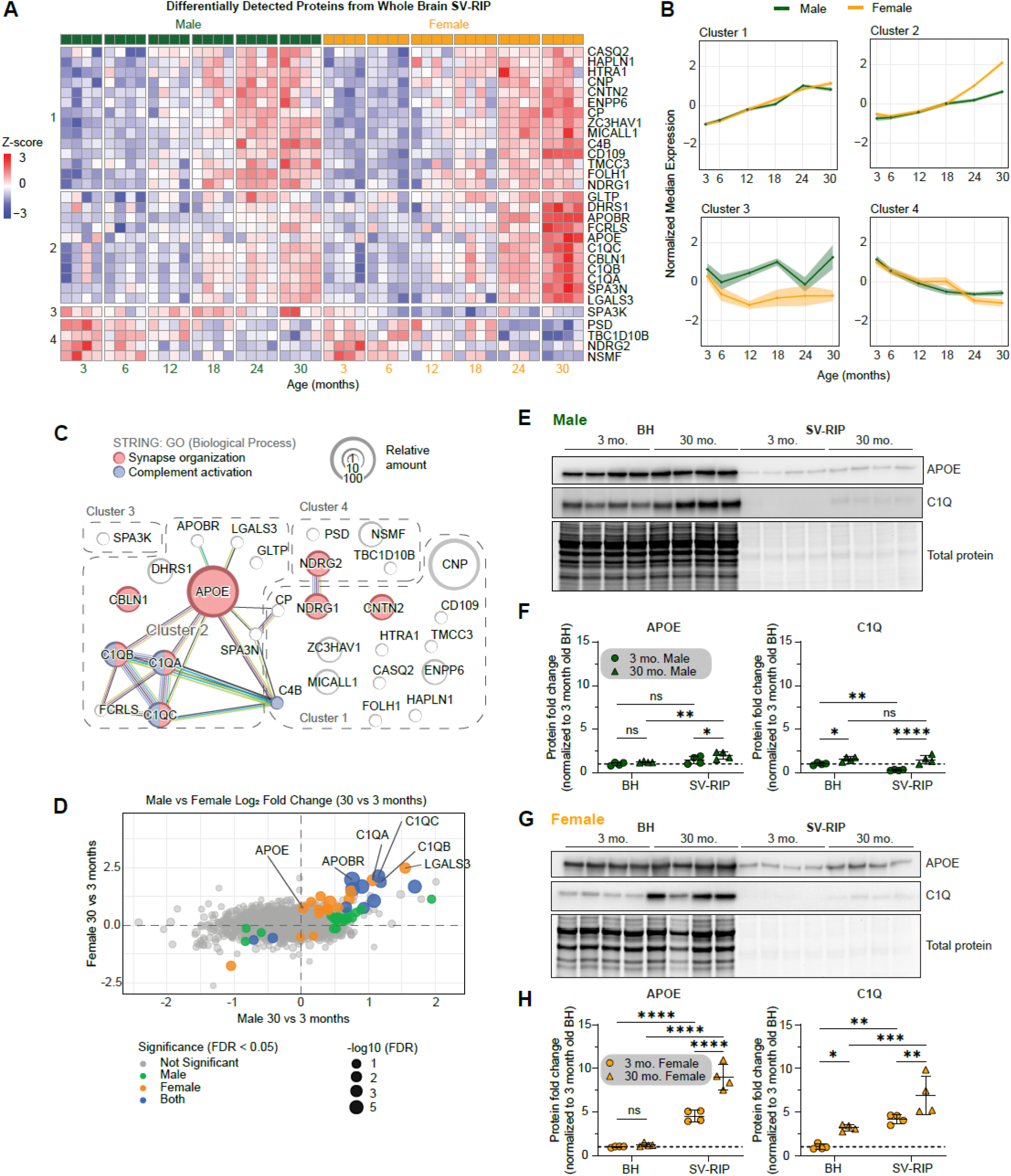
Apolipoprotein E is an abundant synaptic vesicle protein and accumulates with age in females. **A.** Linear regression identified 30 proteins differentially detected (DDPs) across age groups with FDR < 0.05. Proteins were hierarchically clustered using k-means and complexHeatmap R package. **B.** Average normalized median expression of proteins in each cluster from A, split by sex (male;green) (female;yellow). **C.** STRING analysis of the 30 DDPs based on biological process. Synapse organization and complement activation processes are enriched. Size of circles represent relative log2 intensities from proteomics. APOE was the most abundant protein detected. **D.** Log2 fold change between 30 and 3 months for male and female samples. Dot size depicts the -log10 FDR value, and color indicates comparison with significance. **E-F.** Immunoblot analysis of BH and SV-RIP fractions from male samples. APOE and C1Q are mildly increased in SV-RIP. Data are displayed as mean ± SD and analyzed by 2-way ANOVA with Fisher’s LSD test (* p<0.05, ** p<0.01, **** p<0.0001). **G-H.** Immunoblot analysis of BH and SV-RIP fractions from female samples. APOE and C1Q are greatly increased in SV-RIP. Importantly, APOE is not increased in BH fractions, indicating a specific SV interaction. Data are displayed as mean ± SD and analyzed by 2-way ANOVA with Fisher’s LSD test (* p<0.05, ** p<0.01, *** p<0.001, **** p<0.0001).

### Synaptic vesicle lipidomics reveals stable cholesterol with modest phospholipid remodeling across the mouse lifespan in both sexes

Lipids play a key role in SV cycle regulation and impact multiple SV protein interactions **(Binotti, Jahn, & Perez-Lara, 2021)**. We therefore reserved a fraction of our original SV-RIP isolate for untargeted lipidomics and targeted cholesterol analysis (**Figure 4A**). While total protein remained constant throughout age, total phospholipid signal increased in both males and females, suggesting that aged SVs are more lipid-rich, whereas SV cholesterol remained unchanged across age and sex (**Figure 4B-C**). Unsupervised analyses (PCA and sample-sample correlations) showed minimal separation by group (**Supplemental Figure 5A-C**). A heatmap of the top 25 most abundant lipids highlighted a composition dominated by canonical SV lipid species reported previously (red;**(Takamori et al., 2006)**) (**Figure 4D**). Aside from modest age-associated shifts, the most abundant lipid species showed no broad remodeling across the mouse lifespan. We next used LipidMaps BioPAN to infer pathway-level changes from the lipid profiles **(Gaud et al., 2021)**. BioPAN suggested a relative enrichment toward phosphatidic acid (PA) in aged male SVs (3 vs 30 months), consistent with enhanced phospholipid remodeling (e.g., PLD-linked reactions), whereas no pathway reached significance in the analogous female comparison (**Figure 4E–F**). Sex-stratified analyses suggested a bias toward phosphatidylserine (PS) in young female SVs and broader headgroup remodeling in aged females relative to aged males (**Figure 4G-H**). Lipid class and shape-based groupings (cone/cylinder/inverted) and saturation categories showed no significant shifts. All lipid class comparisons are graphed as a percent of total lipid detected in the supplement (**Supplemental Figures 6-7**). Together, these data mirror the proteomic results, the SV lipidome is largely stable across the mouse lifespan, with cholesterol notably stable despite increased SV association of APOE, arguing against a clear lipid-homeostasis role for APOE at the presynapse.

**Figure 4.**
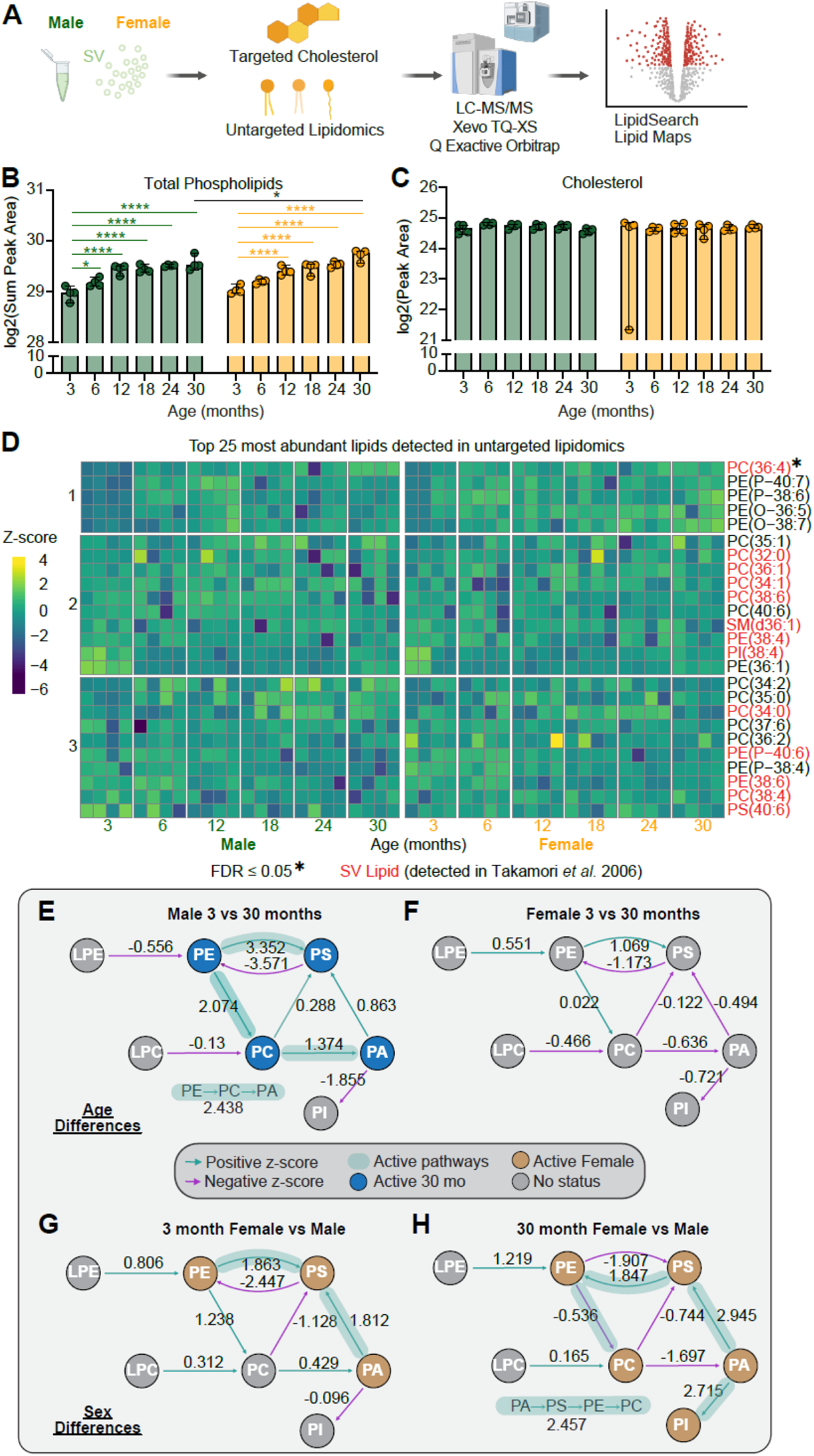
Synaptic vesicle lipidomics reveals stable cholesterol with modest phospholipid remodeling across the mouse lifespan in both sexes. **A.** Experimental workflow for lipidomic analysis of SVs. SVs were isolated from the whole brain of male and female C57BL6/J mice at 3, 6, 12, 18, 24, and 30 months of age by SV-RIP and divided into 4 aliquots (n=4/group, N=48). One aliquot was taken for targeted cholesterol analysis, and another was used for untargeted lipidomics. **B.** Total phospholipids detected with untargeted lipidomics run in positive and negative modes. Data are displayed as means ± SD and analyzed by 2-way ANOVA with Tukey’s multiple comparisons test (* p<0.05, ** p<0.01, ***p<0.001, ****p<0.0001). **C.** Targeted cholesterol detected. Data are displayed as means ± SD and analyzed by 2-way ANOVA with Tukey’s multiple comparisons test (* p<0.05, ** p<0.01, ***p<0.001, ****p<0.0001). **D.** Top 25 most abundant phospholipids detected from untargeted lipidomics, displayed as z-score of percent total phospholipids. SV phospholipids as identified by Takamori et. al, 2006 are labeled in red, and those with an FDR ≤ 0.05 are labeled with an asterisk (limma, multiple comparisons with Benjamini-Hochberg multiple testing correction). **E-H.** BioPAN analysis of detected phospholipids to identify lipid pathways activated/suppressed with age (d, e) and sex (f, g). Active nodes are colored in blue or tan (age and sex differences, respectively), with highlighted arrows representing significantly active/suppressed pathways (z-score ≥ 1.645).

### APOE is broadly enriched at synapses across cortex and hippocampal regions in situ

Because homogenization disrupts native compartmentalization, co-immunoprecipitation (IP) experiments capture any possible interactions, rather than native protein-protein interactions occurring in intact tissue. Therefore, we sought to validate our results with *in situ* localization assays that preserve cellular architecture. To this end, we obtained APOE knockout (KO) mice (JAX:002052) for APOE antibody validation and immunofluorescence-based localization experiments. We compared multiple fixation conditions and found that, although each yielded clear differences between WT and APOE KO tissue, a glyoxal-based fixation recipe produced the lowest background while preserving native structure, and we used this method for all subsequent experiments (**Supplemental Figure 8A, B, D**) **(Richter et al., 2018)**. A coronal section of brain tissue from a WT (C57BL/6J) mouse reveals widespread APOE signal throughout the hippocampus and cortical regions, compared to KO controls (**Figure 5A**). Counterstaining APOE with SYP and DAPI in the sagittal plane further reveals APOE enrichment in the dentate gyrus (DG) of the hippocampal formation (**Figure 5B**). We then explored synaptic localization using high magnification confocal microscopy and quantified colocalization between APOE and SYP. As expected, APOE signal was prominent in astroglia, however focusing within neuropil regions, we observed broad colocalization of APOE with SYP positive synaptic punctae. We investigated cortex (**Figure 5C-D**), hippocampal subregions, the CA1 (**Figure 5E-F**), CA3 (**Figure 5G-H**), and dentate gyrus (DG) (**Figure 5I-J**). All regions showed moderate SYP:APOE colocalization in males and females with Pearson’s correlation coefficient (PCC) of ∼0.3 and Manders’ overlap coefficients (M1 and M2) >80%. All APOE KO slices showed ∼0.1 PCC and ∼0 M1 (M1 defined as fraction of SYP that colocalizes with APOE), indicative of antibody specificity. To benchmark the dynamic range of our assay, we also quantified colocalization of SYP with SYT1, a canonical SV marker expected to strongly overlap with SYP, and with Sortilin, a broadly distributed endolysosomal trafficking protein, also present at the presynapse. **(Nykjaer & Willnow, 2012)**. The SYP:SYT1 measurement yielded a PCC of ∼0.6 and the SYP:sortilin, a PCC of ∼0.2 (**Supplemental Figure 9C**). These findings show that APOE is broadly distributed throughout brain tissue with local enrichment at synapses.

**Figure 5.**
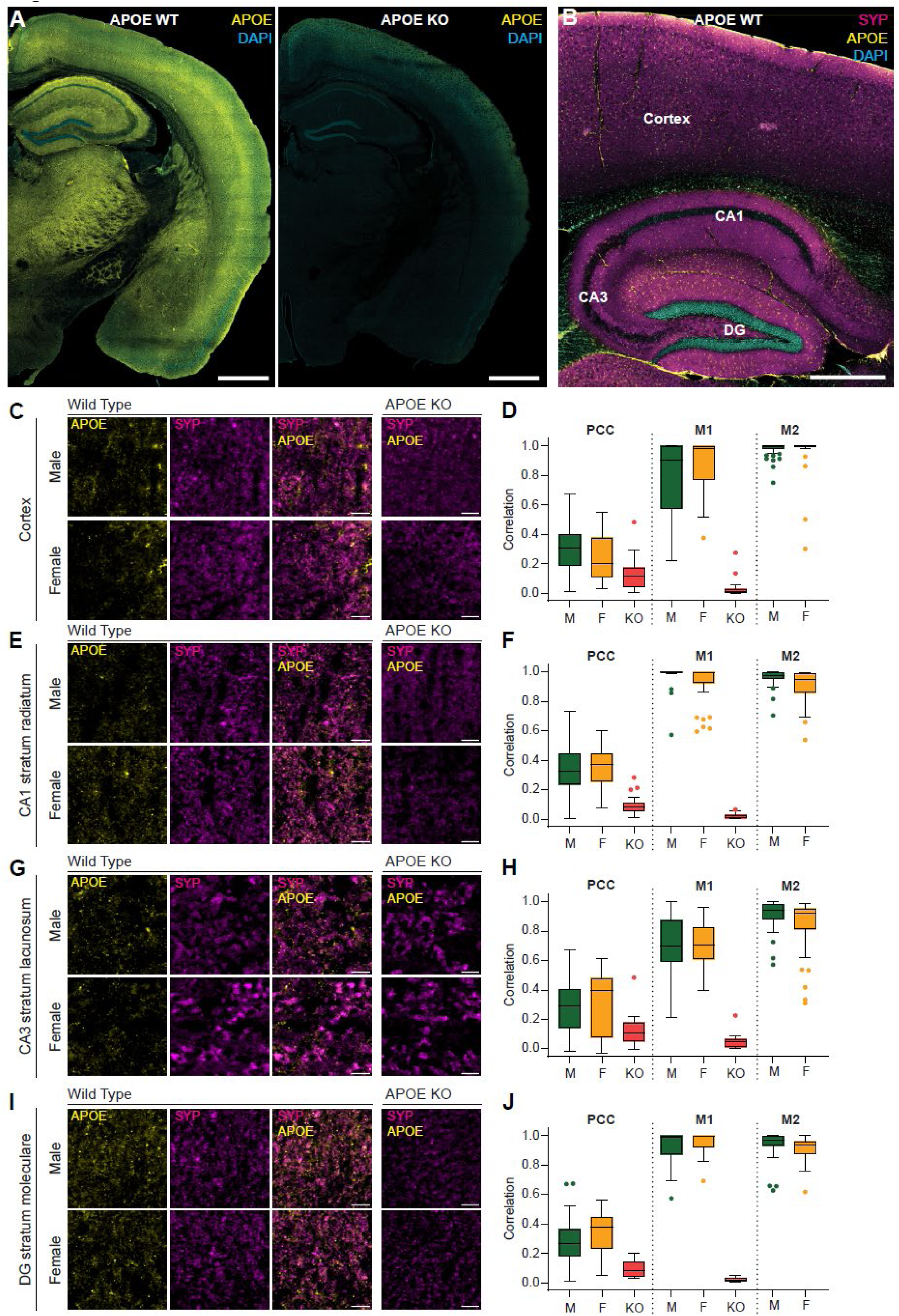
APOE is broadly enriched at synapses across cortex and hippocampal regions in situ. **A.** Coronal section of 3-month-old male mouse brain from APOE WT (C57BL/6J) and APOE KO (B6.129P2-*Apoe^tm1Unc^*/J; JAX#002052) mice, stained with anti-APOE (ab183596) antibody. Scale bar 1 mm. **B.** Sagittal section of a WT mouse brain stained with APOE as in (A) and synaptophysin (SYP; SySy 101308). Cortex, CA1, CA3, and dentate gyrus (DG) labeled. Scale bar 0.5 mm. **C.** Staining of 3-month-old male (top) and female (bottom) cortex with APOE KO controls. Scale bar 5 microns. **D.** APOE KO baseline subtracted measures of colocalization. Pearson’s Correlation Coefficient (PCC), M = male, F = female, KO = APOE KO. Manders Colocalization Coefficients (M1 and M2). M1 measures overlap of SYP with APOE, while M2 measures overlap of APOE with SYP. APOE KO condition is not included with M2 as it is meaningless in a KO. **E.** Staining of 3-month-old male (top) and female (bottom) CA1 region with APOE KO controls. Scale bar 5 microns. **F.** Colocalization quantification as in (D). **G.** Staining of 3-month-old male (top) and female (bottom) CA3 region with APOE KO controls. Scale bar 5 microns. **H.** Colocalization quantification as in (D). **I.** Staining of 3-month-old male (top) and female (bottom) DG region with APOE KO controls. Scale bar 5 microns. **J.** Colocalization quantification as in (D).

### Chemically induced menopause does not alter APOE association with SVs in female mice

Estradiol is neuroprotective and can modulate synaptic function and vulnerability **(Arevalo, Azcoitia, & Garcia-Segura, 2015)**. To test whether ovarian hormone loss contributes to the female biased APOE-SV phenotype, we quantified SV-associated APOE in a chemically induced menopause model. Injection of the ovotoxin 4-vinylcyclohexene diepoxide (VCD) causes gradual follicle depletion, accompanied by decreased estradiol and increased follicle stimulating hormone (FSH), mimicking key features of human menopause **(Brooks, Pollow, & Hoyer, 2016)**. We established a new female C57BL/6J cohort with baseline control groups at 3 and 12 months, and with 12-month mice assigned to vehicle (sesame oil) or VCD treatment and then aged to 18, 24, and 30 months (**Figure 6A**). This experimental design allowed us to 1) repeat our original analysis using aged female controls and now test if loss of hormonal estradiol increased the magnitude of APOE association with SVs. We planned for 10 mice in each condition but through natural attrition, were left with 6-7 mice per group in all but the 30-month cohorts where only 2 survived in each. To confirm VCD-induced ovarian failure, we measured serum FSH before and after injections. We selected FSH due to ease of measurement. Ovarian hormone loss is expected to drive a robust increase in FSH, whereas estradiol often falls toward low levels that can approach assay detection limits. In practice, detecting a rise in an analyte is more reliable than quantifying declines to a near-background level of a hormone. Each animal given VCD had higher FSH two weeks after finishing injections, indicative of ovarian failure, whereas the control groups were unchanged (**Figure 6B**). We performed SV-RIP and TMT-based mass spectrometry on these samples as we did to the original cohort. Again, we found APOE to accumulate with SVs as a function of age starting at 24 months and increasing at 30 months of age in both control and VCD-treated groups (**Figure 6C-D**). We validated the proteomics by immunoblot targeting APOE, SYP, and SYB2 comparing 3-month-old control samples to the 30-month-old samples from the vehicle and VCD treated groups (**Figure 6E**). Confirming the proteomics, we found no change in SYP or SYB2, but increased SV-associated APOE in both the aged vehicle and VCD groups (**Figure 6F**). We conclude that either estradiol does not influence APOE association with SVs, or that VCD treatment initiated at 12 months is too late to reveal such an effect. Furthermore, untargeted lipidomics and targeted cholesterol measurements identified no significantly altered lipids in this cohort (**Figure 6G-H**). Together, these independent data establish APOE as a bona fide age-regulated SV protein and argue against a primary role for accumulated APOE in remodeling the SV lipidome.

**Figure 6:**
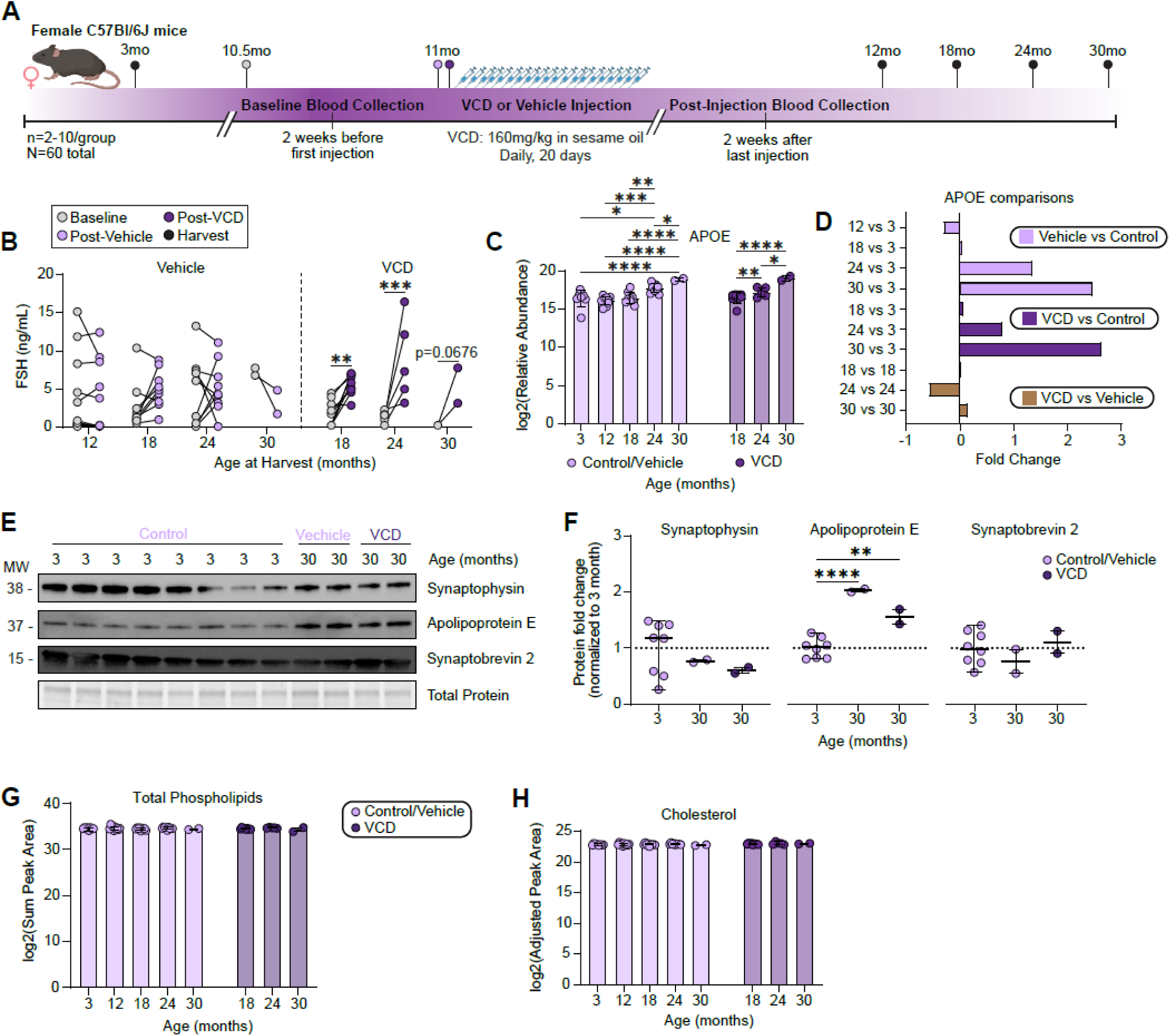
Chemically induced menopause does not alter APOE association with SVs in aged female mice. **A.** Experimental outline for chemically induced ovarian failure using 4-vinylcyclohexene diepoxide (VCD). VCD or vehicle (sesame oil) was injected intraperitoneally for 20 days at 160 mg/kg starting at 11 months of age. Serum samples were collected 2 weeks before and 2 weeks after to assess FSH levels. Mice were aged to 12, 18, 24, and 30 months and SVs isolated from the whole brain with SV-RIP and divided into 4 aliquots. 3-month-old female mice were also collected as young SV and IgG controls (n=2-10/group, N=60 total). One aliquot was taken for TMT proteomics, one for western blot, and one for untargeted lipidomics and targeted cholesterol detection. Only two animals from VCD or CTRL group made it to 30 months in this cohort. **B.** Serum FSH levels (ng/mL) from blood collected at baseline (2 weeks before first injection) and post-VCD or post-vehicle injection (2 weeks after last injection) from female mice collected at 12, 18, 24, and 30 months. Individual values are displayed and matched values connected by a line. Data was analyzed by 2-way repeated measures ANOVA with matching across rows and Tukey’s multiple comparisons test (** p<0.01, *** p<0.001). **C.** Proteomics revealed APOE as the only detected protein that changed in this cohort. Log2 amounts of APOE across cohorts using TMT-based mass spectrometry. **D.** APOE fold change for Vehicle vs Control (Vehicle 12-, 18-, 24-, or 30-month vs Control 3-month), VCD vs Control (VCD 18-, 24-, or 30-month vs Control 3-month), and VCD vs Vehicle (VCD vs Vehicle 18, 24, or 30 month). Importantly, APOE still accumulated in an age dependent manner, confirming our initial analysis. **E.** Immunoblot of Control 3, Vehicle 30, and VCD 30-month SV samples for Synaptophysin, Apolipoprotein E, Synaptobrevin 2, and Total Protein. **F.** Immunoblot quantitation of Control 3, Vehicle 30, and VCD 30-month SV samples, normalized to Control 3 month. Data are displayed as median with 95% confidence intervals and analyzed by 1-way ANOVA with Dunnett’s multiple comparisons test (* p<0.05, ** p<0.01, *** p<0.001, **** p<0.0001). **G.** Total phospholipids detected with untargeted lipidomics run in positive and negative modes. The sum peak area was calculated for all phospholipids within each sample and log2 transformed. Data are displayed as median with 95% confidence intervals and analyzed by 2-way ANOVA with Tukey’s multiple comparisons test. No differences between groups detected. **H.** Targeted cholesterol detected. Peak areas were blank adjusted, IgG subtracted, and log2 transformed. Data are displayed as median with 95% confidence intervals and analyzed by 2-way ANOVA with Tukey’s multiple comparisons test. No differences between groups detected.

### Astroglia-derived APOE is presynaptic and externally oriented on synaptic vesicles

To ensure pre-synaptic localization of APOE, we performed super-resolution imaging on mature dissociated neurons. This approach both validates the culture system for mechanistic studies and localizes APOE at nanometer scale. As APOE is a secreted astroglia protein **(Flowers & Rebeck, 2020)**, so we first verified the presence of GFAP positive cells in our neuronal cultures and found them in abundance in mouse and rat hippocampal, rat cortical, and mouse hippocampal dissociated neuronal cultures, hereafter referred to as co-cultures (**Supplemental Figure 9A-C**) **(Boyles, Pitas, Wilson, Mahley, & Taylor, 1985)**. We then performed immunocytochemistry (ICC) for APOE, SYP and PSD95 on 18 day in vitro (DIV) rat cortical neuron cultures. Restricting our analysis to synapses, we find almost exclusive colocalization of APOE with the presynaptic SYP marker (**Figure 7A-B**). We then used tagged APOE with an astroglia specific promoter to investigate the source of presynaptic APOE. First, we verified the fidelity of the human synapsin (neuron) and GFAP (glia) promoters using cytosolic fluorescent proteins. In rat hippocampal co-cultures co-transduced with 5×10^6^ transducing units (TU) of GFAP: cytosolic mRuby3 and hSyn: cytosolic msGFP lentivirus, we find zero co-expression of these markers and msGFP+ cells have typical neuronal morphology while mRuby3+ cells have typical astroglia morphology (**Figure 7C**). Additional ICC of co-transduced cultures counterstained for anti GFAP confirms astroglia restricted expression of mRuby3 and saturating transduction efficiency in both rat and mouse hippocampal co-cultures (**Supplemental Figure 9D-E**). We then transduced co-cultures with hSyn: SYP-mRuby3 and GFAP: mouse APOE-HaloTag. We confirmed astroglia expression of APOE-HaloTag finding it decorating structures resembling the ER and Golgi (data not shown). APOE generally accepts tags at the carboxy terminus and recently an APOE-HaloTag mouse was created and found to have normal APOE function **(Kaji et al., 2025)**. Co-expression of these constructs along with labeling HaloTag with JF646 **(Grimm et al., 2017)** revealed a high degree of co-localization, confirming that glia-secreted APOE traffics to the presynapse. Finally, we examined SV-associated APOE topology using SVs isolated from whole rat brain by SV-RIP, followed by protease protection analysis (**Figure 7E**). Immunoblotting with a luminal SYT1 antibody and our validated APOE antibody showed that under native conditions, trypsin treatment completely degraded APOE, whereas the luminal SYT1 fragment was protected unless membranes were solubilized with Triton X-100. These data indicate that APOE is on the cytosolic face of the SV membrane (**Figure 7F**). These results reveal APOE is an astroglia-secreted protein that is taken up by neurons and trafficked to the presynapse, interacting with the cytosolic face of SVs.

**Figure 7:**
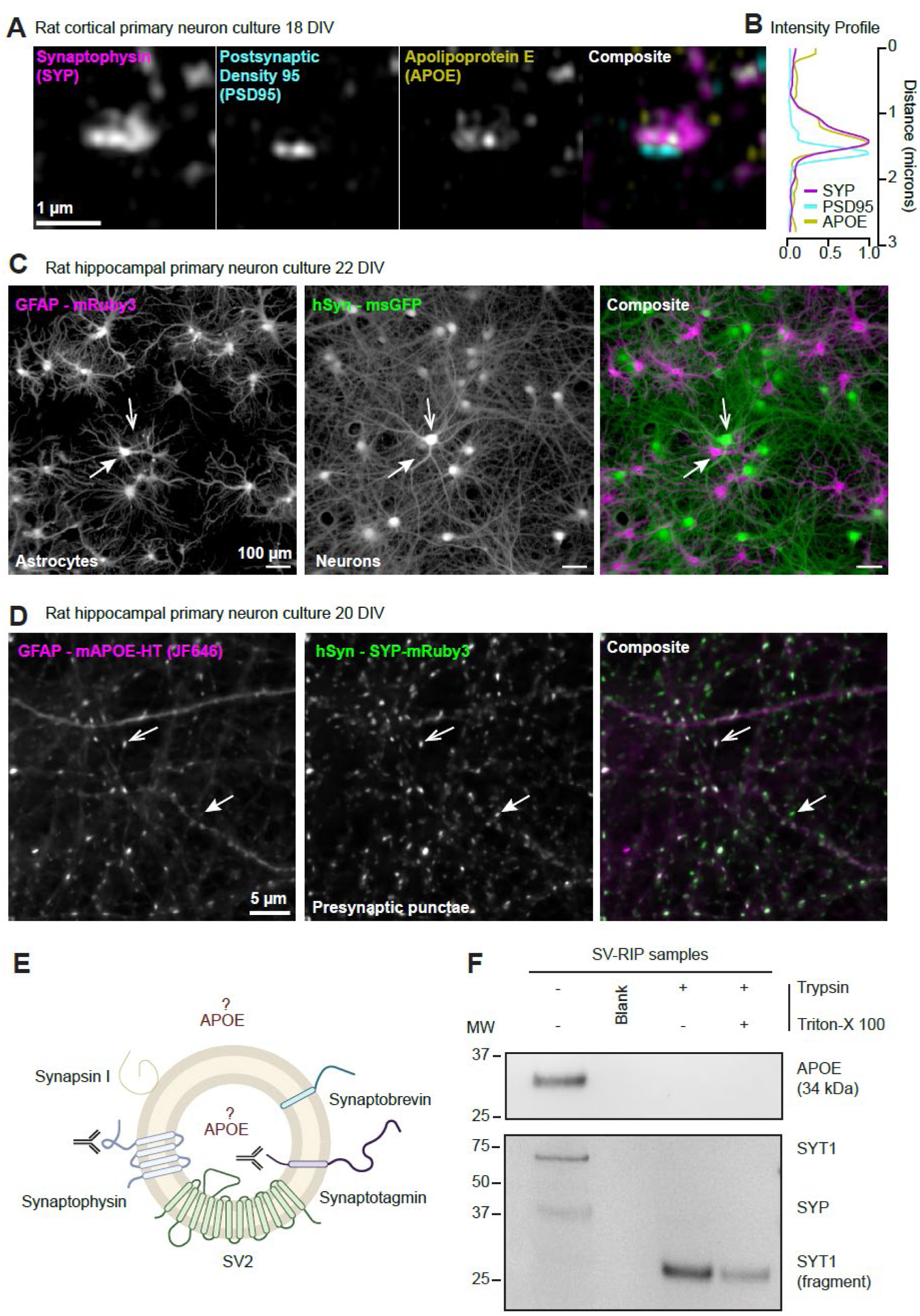
Astrocyte-derived APOE is presynaptic and externally oriented on synaptic vesicles. **A.** Super-resolution imaging via Airyscan confocal and Weiner deconvolution of rat cortical primary neuron-glia cocultures at 18 DIV, stained with antibodies to synaptophysin (magenta), post-synaptic density 95 (cyan), and KO-validated APOE (yellow). Scale bar = 1 micron. **B.** Individual channels were analyzed by line-intensity from top to bottom, normalized to maximum intensity, and plotted together. SYP and APOE signals are distinct from PSD95. **C.** Rat hippocampal primary cocultures transduced with high titers of GFAP-mRuby3 and hSyn-msGFP lentivirus at 5 DIV, and imaged at 22 DIV. Closed arrowhead identifies glia, open arrowhead identifies neuron. **D.** Rat hippocampal primary cocultures transduced with GFAP- mAPOE-HaloTag (mAPOE = mouse APOE) and hSyn – SYP-mRuby3 at 5 DIV and live cell imaged at 20 DIV. HaloTag is localized using HTL-JF646, incubated at 100 nM for 30 minutes just prior to imaging. Open arrowhead identifies a presynapse (via SYP) that has abundant APOE-HT, while closed arrowhead identifies a low APOE-HT presynapse. **E.** SV schematic for protease protection assay with relevant SV proteins illustrated and labeled. APOE is represented as luminal and cytosol facing due to unknown localization. Antibody cartoons indicate antibody epitope site. SYT1 antibody is to the luminal domain of SYT1. **F.** SVs isolated using SV-RIP from whole rat brain were split into groups and treated with nothing (control), trypsin only, or trypsin + Triton X-100. Full length proteins are detected in control conditions, while APOE and SYP disappear with trypsin treatment. SYT1 luminal domain (fragment) is left after trypsin only, while it is further processed after permeabilization with Triton X-100.

### Astroglia expressed Apolipoprotein E regulates the synaptic vesicle cycle

The abundance of APOE at the presynapse prompted us to test whether it regulates the SV cycle. We used the synaptic biosensors iGluSnFR4f and vGlut1-pHluorin, which measure glutamate release and SV pH changes during exocytosis and endocytosis, respectively (**Figure 8A**) **(Aggarwal et al., 2025; Vevea & Chapman, 2020**, **2023)**. These biosensors were expressed in mouse and rat hippocampal co-cultures. Mouse APOE knockout (KO) cocultures were generated from the APOE KO line used throughout this study and were validated for loss of APOE (**Supplemental Figure 10A-B**). Rat cocultures were transduced with the GFAP: APOE-HaloTag lentivirus used in Figure 7 at two doses (High (H) and Low (L)), to model age-related presynaptic APOE accumulation, alongside a GFAP:cytosolic mRuby3 control (CTRL). APOE-HaloTag expression was confirmed by immunoblot (**Supplemental Figure 10C-D**).

**Figure 8:**
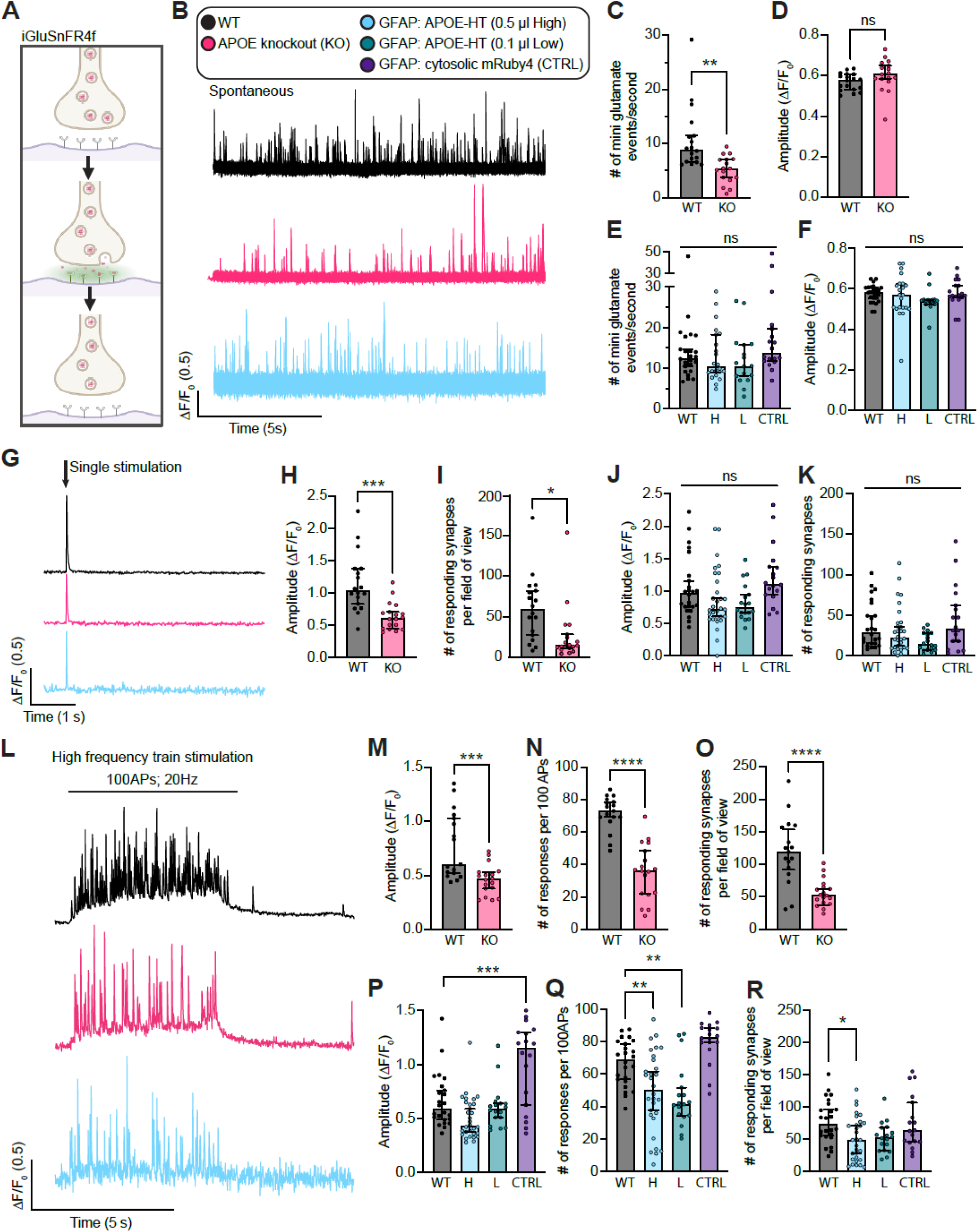
Astrocyte expressed Apolipoprotein E regulates the synaptic vesicle cycle. **A.** Schematic of iGluSnFR4f during exocytosis. **B.** Representative traces of a single region of interest from iGluSnFR4f spontaneous recordings in WT (black), APOE KO (pink) and GFAP: APOE-HT (0.5 µl OE, light blue) hippocampal cultures. **C.** Spontaneous firing frequency of WT and APOE KO hippocampal cultures. Values are median ± 95% confidence interval, n = 17-18 neurons per condition. Welch’s T-test (** p<0.01). **D.** Amplitude of spontaneous events (ΔF/F_0_) for WT and APOE hippocampal cultures. Values are median ± 95% confidence interval, n = 17-18 neurons per condition. Welch’s T-test. **E.** Spontaneous firing frequency of WT, GFAP: APOE-HT (0.5 µl OE), GFAP: APOE-HT (0.1 µl OE) and GFAP: cytosolic mRuby4 (CTRL) hippocampal cultures. APOE-HT was overexpressed using lentiviral transduction (0.5 µl lentiviral volume = High or H, light blue; 0.1 µl = Low or L, teal) of GFAP promoter driven transgene. To control for lentivirus transduction, other cultures were transduced with GFAP promoter driven cytosolic mRuby4 (CTRL, purple). Values are median ± 95% confidence interval, n = 17-22 neurons per condition. One-way ANOVA with Dunnett’s multiple comparisons test. **F.** Amplitude of spontaneous events (ΔF/F_0_) for WT, GFAP: APOE-HT (0.5 µl, H), GFAP: APOE-HT (0.1 µl, L) and GFAP: cytosolic mRuby4 (CTRL) hippocampal cultures. Values are median ± 95% confidence interval, n = 17-22 neurons per condition. One-way ANOVA with Dunnett’s multiple comparisons test. **G.** Single representative traces of iGluSnFR4f single stimulus recording from WT, APOE KO and 0.5 µl (High) GFAP APOE-HT hippocampal cultures. **H.** Amplitude of evoked events (ΔF/F_0_) from WT and APOE KO hippocampal cultures. Values are median ± 95% confidence interval, n = 18 neurons per condition. Welch’s T-test (*** p<0.001). **I.** Number of responding synapses per field of view from WT and APOE KO hippocampal cultures. Values are median ± 95% confidence interval, n = 18 neurons per condition. Welch’s T-test (* p<0.05). **J.** Amplitude of evoked event (ΔF/F_0_) from WT, GFAP: APOE-HT (0.5 µl, H), GFAP: APOE-HT (0.1 µl, L) and GFAP: cytosolic mRuby4 (CTRL) hippocampal cultures. Values are median ± 95% confidence interval, n = 18-30 neurons per condition. One-way ANOVA with Dunnett’s multiple comparisons test. **K.** Number of responding synapses per field of view from WT, GFAP: APOE-HT (0.5 µl, H), GFAP: APOE-HT (0.1 µl, L) and GFAP: cytosolic mRuby4 (CTRL). Values are median ± 95% confidence interval, n = 18-30 neurons per condition. One-way ANOVA with Dunnett’s multiple comparisons test. **L.** Single representative traces of high frequency train stimulus iGluSnFR4f: 100 action potentials; 20Hz train stimulus recording from WT, APOE KO and GFAP: APOE-HT (0.5 µl, H) hippocampal cultures. **M.** Amplitude of evoked events (ΔF/F_0_) from WT and APOE KO hippocampal cultures. Values are median ± 95% confidence interval, n = 17-18 neurons per condition. Welch’s T-test (*** p<0.001). **N.** Number of responses per individual synapse from WT and APOE KO hippocampal cultures. Values are median ± 95% confidence interval, n = 17-18 neurons per condition. Welch’s T-test (**** p<0.0001). **O.** Number of responding synapses per field of view from WT and APOE KO hippocampal cultures. Values are median ± 95% confidence interval, n = 17-18 neurons per condition. Welch’s T-test (**** p<0.0001). **P.** Amplitude of evoked events (ΔF/F_0_) from WT, GFAP: APOE-HT (0.5 µl, H), GFAP: APOE-HT (0.1 µl, L) and GFAP: cytosolic mRuby4 (CTRL) hippocampal cultures. Values are median ± 95% confidence interval, n = 18-30 neurons per condition. One-way ANOVA with Dunnett’s multiple comparisons test (***p<0.001). **Q.** Number of responses per individual synapse from WT, GFAP: APOE-HT (0.5 µl, H), GFAP: APOE-HT (0.1 µl, L) and GFAP: cytosolic mRuby4 (CTRL). Values are median ± 95% confidence interval, n = 18-30 neurons per condition. One-way ANOVA with Dunnett’s multiple comparisons test (**p<0.01). **R.** Number of responding synapses per field of view from WT, GFAP: APOE-HT (0.5 µl, H), GFAP: APOE-HT (0.1 µl, L) and GFAP: cytosolic mRuby4 (CTRL). Values are median ± 95% confidence interval, n = 12-24 neurons per condition. One-way ANOVA with Dunnett’s multiple comparisons test (*p<0.05).

We first measured spontaneous SV release as miniature glutamate transients (mGTs) in the presence of tetrodotoxin (TTX). APOE KO cocultures showed an approximately twofold reduction in mGT frequency, with no change in event amplitude, relative to WT controls (**Figure 8B-D**). Because APOE supports synaptic maturation, we examined synapse structure and found that presynaptic number and size were reduced in APOE KO cocultures, which likely contributes to the spontaneous release defect (**Supplemental Figure 10E-G**). In contrast, astroglia APOE-HaloTag overexpression did not alter mGT frequency at either dose (**Figure 8B, E-F**). We next measured evoked glutamate release in response to a single electrical stimulus. In APOE KO co-cultures, the evoked iGluSnFR4f response was reduced by about half, and the number of responsive synapses per field of view was reduced to roughly one-fifth of control levels (**Figure 8G-I**). Astroglia APOE-HaloTag overexpression again produced little to no effect under these baseline stimulation conditions (**Figure 8G, J-K**).

We then asked whether APOE is required for sustained neurotransmission during high frequency stimulation (HFS). Initial vGlut1-pHluorin experiments showed that both APOE KO and APOE-HaloTag overexpression impaired the ability of synapses to maintain release during HFS (**Supplemental Figure 10H**). APOE KO cocultures could not reliably sustain a 100-action potential train at 20 Hz, so we reduced the stimulus to five trains of 50 action potentials at 10 Hz for KO versus wild-type comparisons. Under these conditions, APOE KO still impaired exocytosis, with little effect on endocytosis (**Supplemental Figure 10I-K**). High-dose APOE-HaloTag overexpression, which increased APOE protein about twofold and approximated the age-dependent increase in SV-associated APOE, also impaired exocytosis and showed a trend toward impaired endocytosis (**Supplemental Figure 10L-N**). To define the exocytic defect more clearly, we returned to iGluSnFR4f and applied 100 action potentials at 20 Hz (**Figure 8L**). APOE KO synapses showed a reduced mean evoked response amplitude (**Figure 8M**). They also failed more often during the train. WT synapses maintained release across the train, with ∼ 60-80% of stimuli eliciting detectable glutamate release, whereas APOE KO synapses released on fewer than 40% of stimuli (**Figure 8N**). Additionally, the number of active synapses per field of view was reduced by about twofold in APOE KO cultures (**Figure 8O**). These data indicate that APOE loss reduces the number of release competent synapses and that the remaining synapses exhibit impaired reliability during sustained stimulation. Strikingly, astroglia-restricted APOE-HaloTag OE produced a related phenotype during HFS. The average amplitude of events (**Figure 8P**), release probability (**Figure 8Q**), and number of active synapses per FOV (**Figure 8R**) were all lower, similar to APOE KO conditions. Altogether, these results establish APOE as a cell non-autonomous regulator of the SV cycle.

## Discussion

Using a rapid whole-organelle immunoprecipitation approach (SV-RIP), we stratified age- and sex-dependent molecular changes in synaptic vesicles (SVs) and identified apolipoprotein E (APOE) as an abundant SV protein that accumulates with age in female mice. We then localized APOE to the presynaptic compartment using intact brain tissue and dissociated primary neuron–astroglia co-cultures. A protease protection assay supports an SV-associated topology consistent with cytosolic exposure. Finally, we assessed presynaptic function with loss of APOE, APOE KO, and astroglia-driven APOE-HaloTag OE in primary neuron co-culture system using iGluSnFR4f and vGLUT1-pHluorin. Using these optophysiology tools, APOE manipulations altered the SV cycle consistent with a modulatory role in SV exocytosis.

We began with discovery-based molecular characterization of the SV using a rapid-immunoprecipitation approach that isolates SVs using the canonical and conserved SV protein SV2 **(Bradberry et al., 2022)**. After validating SV enrichment in our hands, we designed a lifespan, sex-stratified analysis of wild-type C57BL/6J mice at 3-, 6-, 12-, 18-, 24-, and 30-months to identify SV-associated molecular changes that track with the major risk factors for neurodegenerative disease: advanced age and female sex. This design addressed two considerations: first, age-associated synaptic phenotypes emerge gradually, making multi-timepoint sampling well suited to identify factors that change progressively; and second, sex is a major modifier of dementia risk in humans, motivating parallel assessment of male and female trajectories **(Fang et al., 2025; Niccoli & Partridge, 2012)**. In this dataset, we identified a set of proteins with increased SV association as a function of age and sex, with higher levels of these proteins in female samples at older ages. Strikingly, this cluster included proteins implicated in age-related dementia and Alzheimer’s Disease (e.g., APOE and C1q), as well as factors related to innate immune regulation (e.g., LGALS3) and lipid homeostasis (e.g., APOBR) **(Corder et al., 1993; Hong et al., 2016)**. Because the differential list is enriched for non-neuronal secreted proteins, our data suggest that aging may increase SV association of extracellular and immune-related factors that influence synaptic function, motivating direct *in situ* validation of localization and mechanism.

From the 30 differentially detected proteins in Figure 3, APOE stood out based on both disease relevance and high abundance relative to other detected proteins. Indeed, APOE is also among the most prominent proteins in prior SV proteome studies **(Bradberry et al., 2022; Takamori et al., 2006; Taoufiq et al., 2020)**. We document broad APOE synaptic localization in mouse brain tissue and in a reduced and tractable dissociated rodent neuron-astroglia coculture. Using this reduced model, we confirm that astroglia derived APOE accumulates at the presynapse. Independent evidence further links APOE to the synaptic compartment. In human AD brains, APOE localizes to a subset of excitatory synaptosomes using vGLUT1 and NAB61 (oligomeric amyloid-β) and to human synaptosomes prepared using gradient fractionation **(Bilousova et al., 2019; Koffie et al., 2012)**. Additionally, exogenously added human APOE isoforms localize to a fraction of excitatory synapses in dissociated rodent neurons **(Konings, Torres-Garcia, Martinsson, & Gouras, 2021)**. Other reports identified presynaptic proteins syntaxin binding protein 1 and RAB3A as APOE ɛ4 interactors **(Nakamura, Watanabe, Fujino, Hosono, & Michikawa, 2009)**, or report a protein-protein interaction between synaptobrevin 2 (SYB2) and human APOE with a proposed APOE ɛ4 synaptic suppression phenotype **(Chen et al., 2025)**. Most prior localization and functional studies emphasize human APOE isoforms and disease models. While these studies are informative, defining the baseline presynaptic role of APOE in a controlled system is necessary to interpret how isoform- and disease-associated perturbations alter synaptic physiology and pathology.

Here, we provide the first optophysiological analysis of APOE synaptic models using the reporters iGluSnFR4f and vGlut1-pHluorin. These biosensors provide high spatial resolution and capture key features of the SV cycle, including endocytosis, that are not resolved by traditional electrophysiology. Wild-type synapses maintain the SV cycle even at high stimulation frequencies; however, we find that modulating APOE levels lower (APOE KO) or higher (astroglia driven APOE over-expression) suppresses SV exocytosis at high stimulation rates. The defect seems larger in APOE KO as these synapses have defects in spontaneous release, release evoked by single stimulus, and cannot reliably maintain SV exocytosis during HFS at 20 Hz. Notably, our overexpression experiments double APOE expression and increase its accumulation at the presynapse modeling age-related changes identified in this report. APOE is a well-established key mediator of cholesterol transport between glia and neurons, with downstream effects on synaptogenesis and synaptic activity **(Mauch et al., 2001)**. Abundant presynaptic APOE suggests a role in regulating local lipid homeostasis. However, in our aging datasets, SV cholesterol content did not change in aged female samples despite an increased APOE association. A local lipid homeostasis model further predicts that APOE traffics through presynaptic endosomal recycling and resides in the lumen of SVs and presynaptic endosomes. Our protease protection data do not support this, as APOE is protease-accessible and consistent with a cytosolic-facing association. Together, these findings argue against luminal recycling–based lipid regulation as the primary explanation for the presynaptic phenotypes. Our findings motivate an alternative hypothesis that APOE modulates release through a more direct presynaptic mechanism involving protein-protein interactions between APOE and key regulators of SV exocytosis.

APOE has been reported in multiple intracellular compartments. The majority of CNS APOE is produced by astroglia and secreted via the classical ER to Golgi pathway **(Flowers & Rebeck, 2020)**. Lipidated APOE enter neurons via the LDL receptor family gathering in the neuronal endo-lysosomal system where they exchange lipids and recycle back to the extracellular space **(Herz & Bock, 2002; Ioannou et al., 2019; Rensen et al., 2000)** and reviewed in **(Windham & Cohen, 2024)**. Recently, APOE has also been observed to escape ER lumen translocation and traffic to lipid droplets in astroglia **(Windham et al., 2024)**. Other reports also place APOE isoforms with mitochondria and the nucleus, but these reports remain less firmly established and may be dependent on isoform or context **(Chang et al., 2005; Kim et al., 2008)**. This prior work suggests APOE has unusual trafficking behaviors, including avoidance of lysosomal degradation, noncanonical ER processing, and retroendocytic recycling. We identify astroglia-secreted APOE as an abundant SV-associated protein, but the route to this localization is not yet clear. Future work will define the pathway by which APOE reaches the neuronal cytosol and associates with presynaptic compartments. Resolving this trafficking logic is likely to clarify how APOE isoforms modify synaptic vulnerability and neurodegenerative disease risk.

## Resource availability

Data are available upon request from the corresponding author. The mass spectrometry proteomics data have been deposited to the ProteomeXchange Consortium via the PRIDE **(Perez-Riverol et al., 2025)** partner repository with the dataset identifier PXD075522. Lipidomics data are available at Metabolomics Workbench study IDs ST004763 (original aging study) and ST004775 (VCD aging study).

## Acknowledgements

The authors acknowledge and thank the world-class expertise and support provided by The Jackson Laboratory’s Nathan Shock Center of Excellence in the Basic Biology of Aging; animal work was carried out at the center under supervision by Dr. Ron Korstanje and project manager Laura Robinson. We thank Dr. Luke Lavis for HTL-conjugated JF dyes which were graciously provided by his lab and the Janelia Research Campus. We thank Dr. Danielle R. Little and Dr. Michael Dyer for access to APOE knockout animals. We also thank addgene depositors, specifically listed throughout methods. This work would not have been possible without the expert scientists and veterinary technicians throughout St. Jude Children’s Research Hospital including in the Animal Resource Center (ARC), Center for Proteomics and Metabolomics, Vector Laboratory Shared Resource, Biostatistics Shared Resource (BSR), CTIC-EM, and CTIC-LM.

## Author Contributions

Conceptualization, J.D.V.; investigation, S.P., K.B.T., J.S., V.P., G.P., S.D., L.L.R., X.W., B.B., J.C., H.T., A.L., A.J., Z.Y., L.W., C.G.R., A.A.H., J.D.V; methodology, S.P., J.D.V.; resources, funding, R.K., J.D.V.; supervision, J.D.V.; writing - original draft preparation, J.D.V.; writing - review and editing, J.D.V. All authors have read and agreed to the published version of the manuscript.

## Methods

### Ethics statement

Animal care and use in this study were conducted under guidelines set by the National Institutes of Health’s *Guide for the care and use of laboratory animals* handbook. Protocols were reviewed and approved by The Jackson Laboratory’s Institutional Animal Care and Use Committee (AUS#06005) and the Institutional Animal Care and Use Committee of St. Jude Children’s Research Hospital (IACUC protocol #3193).

### Animals

All aged mice used in this study were kindly provided by the Jackson Laboratory Nathan Shock Center. Male and female C57BL/6J mice (C57BL/6J; Strain #:000664; RRID:IMSR_JAX:000664) were included at 3-, 6-, 12-, 18-, 24-, and 30-months of age (n = 4 per age and sex). Animals were maintained under standard conditions in a temperature-controlled facility with a 12-hour light/dark cycle and ad libitum access to regular chow and water. At St. Jude Children’s Research Hospital, time mated Sprague Dawley rats were ordered from Envigo and held for one day before primary neuron cultures were prepared. Wild-type (C57BL/6J; Strain #:000664; RRID:IMSR_JAX:000664) and APOE knockout mice (B6.129P2-Apoetm1Unc/J; JAX strain #:002052; RRID:IMSR_JAX:002052) were maintained under standard conditions in a temperature-controlled facility with a 12-hour light/dark cycle and ad libitum access to regular chow and water.

### SV2a-bead preparation

Dynabeads M-270 epoxy (300 mg, catalog #14302D, Thermo Fisher Scientific) were stored at 4 °C in anhydrous, amine-free N, N-dimethylformamide (DMF) at 30 mg/mL for up to 18 months before coupling. On day 1, 450 µg of either bovine serum IgG (I5506-50MG, Sigma-Aldrich) or anti-SV2α antibody (450 µg; SV2, Genescript Biotech Co., Ltd) was adjusted to 300 µl with 100 mM borate buffer (100 mM boric acid, 76 mM NaOH, pH 8.5). Fifteen milligrams of beads were transferred to a fresh polypropylene tube, DMF was removed on a magnetic stand, and beads were washed once with ice-cold borate buffer. Beads were resuspended in 300 µl borate buffer, mixed with the antibody solution, and combined with 300 µl of 3 M ammonium sulfate in borate buffer. The mixture was incubated overnight at 37 °C with rotation. On day 2, antibody-beads were collected on a magnetic stand, washed alternately with wash buffer I (500 mM NaCl, 50 mM ammonium acetate, pH 4.5) and wash buffer II (500 mM NaCl, 50 mM Tris-HCl, pH 8.0) for three cycles, and resuspended in ice-cold KPBS (145 mM KCl, 10 mM potassium phosphate consisting of KH₂PO₄ and K₂HPO₄, pH 7.4). After transferring to a new tube and a final wash, beads were stored at 4 °C until use.

### Tissue homogenization

The whole mouse brain (including cerebrum, cerebellum and brainstem) was removed and transferred to ice-cold Hibernate™-A medium (A1247501, Gibco). Tissue was homogenized manually in hypotonic buffer (25 mM KCl, 20 mM potassium phosphate buffer consisting of KH₂PO₄ and K₂HPO₄, 5 mM EGTA, pH 7.3) containing protease inhibitors (cOmplete Mini EDTA-free, 1 tablet/10 mL)] using a Teflon-glass Dounce homogenizer. Fifteen strokes were manually applied to the tissue for complete homogenization. The tissue lysate was then centrifuged (20 min, 25,000 × g, 2 °C). Supernatant was collected and adjusted to 125 mM KCl by adding 4 M KCl prior to rapid immunoprecipitation.

### Rapid immunoprecipitation of synaptic vesicles

For each half brain, 7.5 mg of antibody-conjugated beads were used. KPBS was removed from the beads, followed by washing once with 1 mL ice-cold KPBS by vertexing. Fifteen milligrams of SV2α-beads were divided into two 2 mL microcentrifuge tubes and resuspended in 100 µl ice-cold homogenization buffer. To each tube, 1.9 mL of supernatant from the previous step (about 2.5 mg/mL protein, representing vesicles from half a brain) was added. The tubes were placed in 50 mL conical tubes packed with ice and incubated with rotation for 25 minutes in a cold room. Following incubation, beads were separated on a magnetic stand, and the supernatant was transferred to new tubes. Beads carrying synaptic vesicles were washed gently by flipping the tube three times with 1 mL ice-cold KPBS, then resuspended in 0.5 mL fresh ice-cold KPBS. Beads from the same brain were combined into fresh 2 mL microcentrifuge tubes. The bead bound synaptic vesicles were aliquoted into 4 tubes. Each aliquot will be then processed for elusion according to downstream application requirements.

### Negative stain transmission electron microscopy

Freshly glow discharged carbon coated 200 mesh copper grids floated on 10µl drops of sample and absorbed for 10 minutes. The excess sample was removed by carefully wicking fluid from grid surface such that the surface never dried, and grids were washed three times on droplets of milliQ purified water, again wicking to remove excess fluid. Grids were contrasted by floating on drops of 1% uranyl acetate. The excess stain was removed by careful wicking to leave a thin film of stain on the grid surface. Grids were air dried and examined in a ThermoScientific F20 Tecnai transmission electron microscope at 80kV and images captured on an AMT NanoSprint 15 MkII imaging system. All materials and reagents from Electron Microscopy Sciences (Hatfield, PA) unless otherwise stated.

### Negative Stain EM and SV-RIP vesicle analysis

Negative stain EM images were analyzed using Fiji for ImageJ **(Schindelin et al., 2012)** and the SAMJ plugin (Garcia-Lopez-de-Haro et al., 2025).

### In-gel digestion protocol, LC-MS, and spectral counting-based proteomic data analysis

Protein samples were run on a short gel as described in a previously published protocol **(Xu, Duong, & Peng, 2009)**. Protein in gel bands were reduced with dithiothreitol (DTT) (Sigma) and alkylated by iodoacetamide (IAA) (Sigma). The gel bands were then washed, dried, and rehydrated with a buffer containing trypsin (Promega). Samples were digested overnight, acidified and the peptides were extracted. The extracts were dried and reconstituted in 5% Formic acid. The peptide samples were loaded on a nanoscale capillary reverse phase C18 column (Thermo Easy Spray (ES904) 75id, 2um C18, 150mm) by a HPLC system (Thermo Ultimate3000) and eluted by an acetonitrile gradient. The eluted peptides were ionized and detected by an inline mass spectrometer (Thermo Orbitrap Exploris 480). The MS and MS/MS spectra were collected over a 75-min liquid chromatography gradient. The mass spectrometer is operated in data-dependent mode with a survey scan in Orbitrap (60,000 resolution, 1 × 10^6^ AGC target and 50 ms maximal ion time) and MS/MS high resolution scans (15,000 resolution, 1 × 10^5^ AGC target, 150 ms maximal ion time, 32 HCD normalized collision energy, 1.5 *m/z* isolation window, and 8 s dynamic exclusion).

Database searches were performed using JUMP **(X. Wang et al., 2014)** search engine against a composite target / decoy Uniprot mouse protein database **(Elias & Gygi, 2007)**. Searches were performed using a 25-ppm mass tolerance for precursor and fragment ions, fully tryptic restriction with two maximal missed cleavages. Carbamidomethylation of Cysteine (+57.02146 Da) was used for static modification and Met oxidation (+15.99492 Da) was considered as a dynamic modification. All matched MS/MS spectra were filtered by mass accuracy and matching scores to reduce protein false discovery rate to <1%. Spectral counts, matching to individual proteins reflect their relative abundance in one sample after the protein size is normalized. The spectral counts between samples for a given protein is used to calculate the p-value which is derived by G-test **(Zhou et al., 2010)**.

### In-gel digestion protocol, LC-MS, and TMT-based proteomic data analysis

Protein samples were run on a short gel as described in a previously published protocol **(Xu et al., 2009)**. Gel bands were excised and washed twice with 50% acetonitrile and once with 50% acetonitrile in 5 mM HEPES. The gel bands were then dried in a speed vacuum. The dried gel plugs were rehydrated with a buffer containing trypsin in 5 mM HEPES (Promega). The samples were digested overnight at 37^0^C. The peptides from the gel plugs were extracted with 100% acetonitrile. The peptide extracts were then dried down and resuspended in 50 mM HEPES (pH 8.5). The peptides were labeled with 18-plex Tandem Mass Tag (TMT) reagents (Thermo Scientific) following the manufacturer’s recommendations. After labeling with TMT reagents for 1 hour, the unreacted labels were quenched with 5% hydroxylamine. The TMT-labeled samples were mixed equally and desalted using C18 cartridges (Harvard Apparatus). The peptide eluent from the cartridge was dried in a speed vacuum and resuspended in Buffer A for basic pH reverse-phase liquid chromatography. The sample was then fractionated on an offline HPLC (Agilent 1220) using basic pH reverse-phase liquid chromatography (pH 8.0, Waters XBridge C18 column, 2.1 mm × 10 cm, 3.5 µm particle size). For basic reverse phase fractionation, buffer A consisted of 10 mM NH_4_COOH (pH 8.0) whereas buffer B consisted of 10 mM NH_4_COOH (pH 8.0) / 90% acetonitrile. 200 µl/min flow rate was used, and 60 one-minute fractions were collected and concatenated back to 30 fractions (i.e., the 1^st^ fraction was pooled with the 31^st^ fraction, the 2^nd^ fraction with 32^nd^ fraction, and so on). The fractions were dried and resuspended in 5% formic acid and analyzed by acidic pH reverse phase LC-MS/MS. For acidic pH reverse phase LC-MS/MS analysis, buffer A consisted of 0.2% formic acid whereas buffer B was 70% acetonitrile in 0.2% formic acid. The peptide samples were loaded on a nanoscale capillary reverse phase C18 column (Thermo Easy Spray (ES904) 75id, 2um C18, 150mm) using an HPLC system (Thermo Ultimate 3000) and eluted with a 95-min gradient from 5%B-95%B at 350 nL/min. The eluted peptides were ionized by electrospray ionization and detected by an inline Orbitrap Exploris 480 (Thermo Scientific) mass spectrometer. The mass spectrometer was operated in a data-dependent mode with a MS scan in the Orbitrap (60,000 resolution, and 50 ms maximal ion time) followed by MS/MS high-resolution scans (60,000 resolution,105 ms maximal ion time, 34 HCD normalized collision energy, 1 *m/z* isolation window). A 15 second dynamic exclusion was used during data acquisition.

All the MS data were processed with an in-house JUMP software suite, a tag-based hybrid search engine to improve sensitivity **(X. Wang et al., 2014)**. The protein database was downloaded from uniport.org (Swiss-Prot and TrEMBL) in January of 2015 (52,738 entries for mouse proteins), followed by concatenation with a decoy database. Major parameters included 15 ppm mass tolerance for precursor ions and 15 ppm for product ions, full trypticity, static modification of the TMTpro tags (+304.20714 Da) on Lys residues and peptide N termini, dynamic modification for Met oxidation (+15.99492 Da), maximal miscleavage sites (*n* = 2), and maximal modification sites (*n* = 3). The resulting PSMs are filtered to reduce protein FDR below 1%. The protein quantification was based on TMT reporter ion intensities with *y*_1_-ion based correction of TMT data to reduce the effect of ratio compression **(Niu et al., 2017)**. Normalization method was using internal standard for batch effect removal on 3 batches by an in-house program JUMP Batch. The average IgG intensity was subtracted from other samples for each protein. Differential protein expression analysis was done by the in-house program JUMPshiny **(Zhang et al., 2025)**. Co-expression clusters were identified, and pathway enrichment was done by the in-house JUMPn software **(Vanderwall et al., 2021)**.

### Lipid extraction

An aliquot of synaptic vesicles bound to SV2-conjugated Dynabeads were flash frozen and stored at −80 °C until lipid extraction. Total lipids were extracted using a modified Folch extraction procedure as previously described **(Boada-Romero et al., 2025; Folch, Lees, & Sloane Stanley, 1957)**. Briefly, SV-beads were resuspended in 0.2mL of ice-cold 0.9% NaCl. 1mL chloroform-methanol (2:1, vol/vol) was added to the samples and mixed by vortexing. The mixture was homogenized for 30 seconds at 4 m s^-1^ using a Bead Ruptor Elite (OMNI International, Kennesaw, GA, USA). Samples were then centrifuged for 10 minutes at 21,000g at 4 °C. The lower organic phase layer was carefully transferred to a new tube and allowed to evaporate to dryness under a stream of liquid nitrogen. The dried lipid extracts were thoroughly redissolved in 50 µl chloroform-methanol (2:1, vol/vol), transferred to autosampler vials and analyzed by lipid chromatography with MS/MS for untargeted lipidomics and targeted cholesterol analysis (LC-MS/MS; 10 µl per injection for untargeted lipidomics, 4 µl per injection for targeted cholesterol).

### LC-MS/MS lipid profiling

#### Untargeted lipidomics

Liquid chromatography separation was performed using a Vanquish Horizon UHPLC system (ThermoFisher Scientific) with the following stepped-gradient conditions: 0-4.5 mins, 45-60% mobile phase B; 4.5-5.0 min, 60-70% B; 5-8 min, 70% B; 8-19 min, 70-75% B; 19-20 min, 75-90%, B; 20-33 min, 90-95% B; 33-34 min, 90-100% B; 34-39 min, 100% B; 39-40 min, 100 to 45% B; 40-45 min, 45% B. Mobile phase A was water/acetonitrile (60:40, vol/vol) and mobile phase B was isopropanol/acetonitrile (90:10, vol/vol); both contained 10 mM ammonium acetate. The column used was a ThermoFisher Scientific Accucore C30 (2.1 mm x 250 mm, 2.6 µm) operated at 50 °C. The flow rate was 250 µl/min and the injection volume was 10 µl. The analysis was conducted using a ThermoFisher Scientific Q Exactive hybrid quadropole-Orbitrap mass spectrometer (QE-MS) equipped with a HESI-II probe as the detector. Two chromatographic runs were performed for each sample, acquiring data separately for both negative and positive ions. The QE-MS was operated using a data-dependent LC-MS/MS mode (Top-15 dd-MS^2^) for both ion modes. The mass spectrometer was set at a resolution of 140,000 (full width at half maximum at *m/z* 200), and AGC target of 1 × 10^6^ and maximum injection time of 80 ms. The scanning range was 100-1,500 *m/z*. The instrument’s operating conditions were as follows: sheath gas flow, 45; aux gas flow, 8; sweep gas, 2; spray voltage, 3.5 kV (positive mode) and 2.5 kV (negative mode); capillary temperature, 320 °C; S-lenses RF level 50; and aux gas heater temperature, 320 °C. For the Top-15 dd-MS^2^ conditions, the resolution was set to 35,000, AGC target of 1 × 10^5^, maximum injection time of 50 ms, MS^2^ isolation width of 1.0 *m/z* and normalized collision energy (NCE) of 35.

#### Targeted cholesterol

An authentic cholesterol standard was purchased from Sigma-Aldrich. Liquid chromatography separation was performed using a Xevo triple quadropole mass spectrometer (Waters) with 100% mobile phase B comprised of isopropanol/acetonitrile (90:10, vol/vol) with 10 mM ammonium acetate. The column used was a Phenomenex kinetex Biphenyl (50 mm x 2.1 mm; 1.7 µm) operated at 45 °C. The flow rate was 400 µl/min and the injection volume was 4 µl. The analysis was conducted using a Waters Xevo triple quadropole mass spectrometer (TQ-MS) equipped with an atmospheric pressure chemical ionization (APCI) probe as the detector. The TQ-MS was operated in positive mode, using a multiple reaction monitoring method (MRM). The APCI source operating conditions were: corona current 10 µA; cone voltage 8 V; desolvation gas flow 1000 L/hr; cone gas flow 150 L/hr; nebulilser gas flow 7 bar; collision gas flow 0.15 ml/min; APCI probe temperature 350 °C; Source temperature 150 °C. The ions targeted (parent/daughter m/z) were 369.43>147.01, 369.43>109.04, 369.43>94.97 and 369.43>80.97 for cholesterol analysis.

### Lipid identification and data analysis

Lipid identification and relative quantification were performed using the ThermoFisher Scientific LipidSearch software (version 5.0) with the following parameters: precursor and product ion mass tolerance, ±5 ppm; and main adducts search, M+H, M-H, M+NH_4_, M+CH_3_COO, M+2H, M-2H, M+Na, M+K. Lipid subclasses – including phospholipids, sphingolipids, glycerolipids and neutral lipids – were searched for the major lipid classes. All individual data files were analyzed for product ion MS/MS spectra of lipid precursor ions. The MS/MS predicted fragment ions for each precursor adduct were measured within 5 ppm of mass tolerance. Product ions matching the predicted fragment ions within this tolerance were used to calculate a match score, with the highest quality matches being selected for lipid molecule identification. Peak areas were integrated to generate chromatographic data for semiquantitative analyses. Following lipid identification, alignment of the resulting data across the experimental groups was performed with the following alignment parameters: experiment type, LC-MS; alignment method, mean; retention time tolerance, 0.25 min; calculate unassigned peak area, on; filter type, new filter; toprank filter, on; main node filter, all isomer peaks; m-score threshold, 5.0; c-score threshold, 2.0; and ID quality filter, A,B,C and D. The sum of all peak areas for each sample was considered as total lipid content. The LipidSearch results were exported to Excel and the Area column used for further analysis. First, the average IgG value was subtracted from the sample values for each lipid, and lipids with the same sum composition ([Lipid Class] [Total Carbons] : [Total Double Bonds]) were combined by addition of their peak areas. After combining lipids by sum composition, those with negative values for one or more samples were removed and individual lipids normalized to total lipid content to account for variations in lipid extraction or data acquisition. Differential analysis was performed using the in-house JUMPshiny software as was done for the proteomic data.

### Tissue Fixation, Sectioning, and Immunofluorescence

Brains were fixed using a glyoxal-based fixative prepared fresh by mixing 2.835 mL ddH₂O, 0.789 mL absolute ethanol, 0.313 mL glyoxal (40% stock; Sigma-Aldrich #128465), and 0.03 mL acetic acid, and adjusting the pH to ∼4.5 with 5 N NaOH to obtain a final solution containing 3% glyoxal, 0.8% acetic acid, and 20% ethanol **(Richter et al., 2018)**. Mice were deeply anesthetized with pentobarbital (100 mg/kg, intraperitoneally) and transcardially perfused with saline followed by glyoxal fixative (two times the body weight in volume, e.g., ∼60 mL for a 30 g mouse). Brains were removed and post-fixed overnight in the same fixative at 4 °C. Coronal sections (50 µm) were collected using a vibratome (VT1000S, Leica Microsystems) and stored free-floating in 1x PBS containing 0.1% Azide at 4°C. For immunofluorescence, sections were washed in 1x PBS containing 0.1% Triton X-100 (PBS-T), blocked for 60 minutes in 10% normal donkey serum, and incubated with were used at 1:400 dilution. Sections were washed again on next day for three times with PBS-T and mounted onto glass slides for imaging. Images were acquired using a 63× oil immersion objective on a ZEISS LSM 780 laser scanning microscope (ZEISS, Germany) and analyzed with ImageJ software version 1.54p (NIH, Bethesda, MD, USA) **(Schindelin et al., 2012)**.

### Proteinase protection assay (PPA) on synaptic vesicle-bound beads

To examine the localization of APOE on synaptic vesicles, 0.625 mg SV-bead samples were transferred to separate tubes to perform PPA. Samples were incubated directly with the indicated reagents in 1× PBS, pH 7.4. The 3 conditions were: 1) control, 30 μl PBS; 2) trypsin treatment, 29 μl PBS plus 1 μl trypsin (1 mg/ml working concentration); and 3) detergent-permeabilized trypsin treatment, 24.5 μl PBS plus 1 μl trypsin (1 mg/ml) and 4.5 μl 2% Triton X-100. Samples were incubated for 10 min at 37°C. Reactions were then supplemented with either 10 μl PBS (control) or 5 μl PBS plus 5 μl trypsin inhibitor (1 mg/ml) (trypsin and Triton X-100 conditions) and incubated for an additional 30 min at 37°C. The beads were separated on magnetic stand, and the cleared supernatant was transferred to a new tube. As for positive control, the SV-beads were directly eluted in 40 μl elution buffer (2% SDS and 25 mM Tris-HCl, pH 8.0). All four samples were then analyzed by SDS–PAGE and western blotting.

### Primary neuron cocultures

Mouse hippocampal neurons were isolated from postnatal day 0 mice. Briefly, hippocampal tissue was dissected and maintained in chilled hibernate A media (Gibco; 1247501). This tissue was then incubated with 0.25% trypsin (Corning; 25-053 Cl) for 30 min at 37°C before trituration in DMEM (Gibco; 11965-118) supplemented with 10% fetal bovine serum (FBS, Thermofisher; 501527079) and penicillin-streptomycin (pen/strep, Thermofisher; MT-30-001-Cl). Dissociated neurons were plated on 18 mm glass coverslips (Warner instruments; 64-0734 [CS-18R17]) coated with poly-D-lysine (Thermofisher; ICN10269491) and laminin (Thermofisher; 23017015). Neurons were initially plated in DMEM with 10% FBS and pen/strep for 2 hours at a density of 250,000 cells/well. Then, media was replaced with Neurobasal-A medium (Thermofisher; 10888-022) supplemented with 2% B-27 (Thermofisher; 17504001), 2 mM Glutamax (Thermofisher; 35050031), and pen/strep. Rat hippocampal and cortical neurons were isolated similarly with the following changes. Time mated rats (Sprague Dawley) hippocampal and cortical neurons were isolated from E18 pups of both sexes for primary neuron cocultures. Rat primary cells were plated at a density of 125,000 cells/well.

### Immunocytochemistry

ICC on primary neuron-astroglia cocultures was performed as previously described **(Vevea et al., 2021)**. Primary antibodies used were anti-APOE (1:100) (Abcam; ab183596; RRID:AB_2832971) for Fig. 5A-C, E, G, and I; Fig. 7A; Sup. Fig. 8A and D; and Sup. Fig. 10B, anti-SYP1 (1:100) (Synaptic Systems; 101 004; RRID:AB_1210382) for Fig. 5; Fig. 7A; Sup. Fig. 8D; and Sup. Fig. 10E, anti-MAP2 (1:100) (Sigma-Aldrich; M9942; RRID:AB_477256) for Sup. Fig. 9A-C and Sup. Fig. 10E, anti-GFAP (1:100) (Agilent; GA52461-2; RRID:AB_2811722) for Sup. Fig. 9A-C, anti-GFAP (1:100) (DSHB; N206A/8; RRID:AB_2877343) for Sup. Fig. 10B, and anti-PSD95 (1:100) (DSHB; K28/43; RRID:AB_2877189) for Fig. 7A. Secondary antibodies used were goat anti-rabbit Alexa Fluor Plus 488 (1:1000) (Thermo Fisher Scientific; A48282TR; RRID:AB_2896346) for Fig. 5A-C, E, G, and I; Fig. 7A; Sup. Fig. 8A and D; Sup. Fig. 9A-C; and Sup. Fig. 10B, goat anti-guinea pig Alexa Fluor 647 (1:1000) (Jackson ImmunoResearch Labs; 106-606-003; RRID:AB_2337449) for Fig. 5B, C, E, G, and I; Fig. 7A; Sup. Fig. 8A and D; and Sup. Fig. 10E, goat anti-mouse Alexa Fluor Plus 555 (1:1000) (Thermo Fisher Scientific; A48287; RRID:AB_2896353) for Fig. 7A, goat anti-mouse Alexa Fluor Plus 647 (1:1000) (Thermo Fisher Scientific; A48289; RRID:AB_2896355) for Sup. Fig. 9A-C and Sup. Fig. 10B, and goat anti-mouse Alexa Fluor Plus 488 (1:1000) (Thermo Fisher Scientific; A48286; RRID:AB_2896351) for Sup. Fig. 10E. Images were acquired using a ZEISS LSM 980 Airyscan laser scanning microscope (ZEISS, Germany) and analyzed with ImageJ software version 1.54p (NIH, Bethesda, MD, USA) **(Schindelin et al., 2012)**.

### Immunoblot

For immunoblots related to SV-RIP, 3.75 mg of SV-bound beads were eluted in 30 µL of elution buffer containing 2% SDS and 25 mM Tris-HCl, pH 8.0. To load the gel, 5 µL of brain homogenate was used as total input and 2 µL of SV eluate was loaded. For immunoblots from primary neuron-astroglia cocultures, each well was lysed directly in sample buffer. Protein concentrations were not measured for primary culture lysates because cells were plated at equal density and lysed in equal volumes; equal loading was confirmed by total protein staining. Samples were combined with XT sample buffer (Bio-Rad, Cat# 1610791) and XT reducing agent (Bio-Rad, Cat# 1610792), heated to 72 °C for 10 min, separated on 18-well 4-12% Criterion XT Bis-Tris gels (Bio-Rad, Cat# 3450124), and transferred to 0.45 µm PVDF membranes (Millipore, Cat# IPFL00010). Following transfer, gels were stained with GelCode Blue Stain Reagent (Thermo Scientific, Cat# 24592) for 1 h, washed in ddH_2_O overnight, and imaged with a 60 s exposure to assess total protein. Membranes were blocked in 5% nonfat milk in TBS-T for 30 min, incubated with primary antibodies overnight, washed three times for 5 min each in TBS-T, and then incubated with HRP-conjugated secondary antibodies. Primary antibodies used were anti-LAMP1 (1:1000; Abcam; ab24170; RRID:AB_775978), anti-VDAC1 (1:1000; Cell Signaling Technology; 4661; RRID:AB_10557420), anti-EEA1 (1:1000; Abcam; ab2900; RRID:AB_2262056), anti-Rab7 (1:1000; Abcam; ab137029; RRID:AB_2629474), anti-Calreticulin (1:1000; Abcam; ab2907; RRID:AB_303402), anti-ATP1A1 (1:1000; Abcam; ab7671; RRID:AB_306023), anti-M6PR (1:1000; Abcam; ab2733; RRID:AB_2122792), anti-sortilin (1:1000; Abcam; ab16640; RRID:AB_2192606), anti-APOE (1:1500; Abcam; ab183596; RRID:AB_2832971), anti-SYT1 (1:1000; DSHB; mAb 48; RRID:AB_2199314), anti-SYB2/VAMP2 (1:1000; Synaptic Systems; 104 211; RRID:AB_887811), anti-SYP1 (1:1000; Synaptic Systems; 101 004; RRID:AB_1210382), anti-PSD95 (1:1000; DSHB; K28/43; RRID:AB_2877189), anti-C1QA (1:1000; Abcam; ab182451; RRID:AB_2732849), and anti-SYT1 luminal domain (1:1000; Synaptic Systems; 105 104; RRID:AB_1106796). Secondary antibodies used were goat anti-rabbit IgG-HRP (1:5000; Bio-Rad; 170-5046; RRID:AB_11125757), goat anti-mouse IgG-HRP (1:5000; Bio-Rad; 170-5047; RRID:AB_11125753), and goat anti-guinea pig IgG-HRP (1:5000; Thermo Fisher Scientific; A18769; RRID:AB_2535546). The immunoblots were imaged using Western HRP Substrate (Millipore Forte, Cat#WBLUF0500) and Biorad ChemiDoc MP Imaging System.

The immunoreactive bands were analyzed using Fiji and the contrast was linearly adjusted for publication. For the core SV proteins in figure 2H-K, each sample was internalized by its total protein prior to its normalization with 3-month-old male to obtain the relative fold change of SYP, SYT1, and SYB2. For figure 3E-F, each sample was internalized by its total protein and normalized to the average of the 3-month-old males and females brain homogenate (BH) respectively, to obtain the relative protein fold change of APOE and C1Q.

### Plasmids

Plasmids used in this study were either purchased from Addgene, or made in house derived from a lentiviral human synapsin modified FUGW backbone ((FUGW was a gift from David Baltimore (Addgene plasmid # 14883; 14883; http://n2t.net/addgene: RRID:Addgene_14883)). We leveraged a fourth generation iGluSnFR (4f) glutamate probe subcloned into our hSynapsin promoter FUGW backbone (pAAV-CAG-iGluSnFR4f-NGR-WPRE was a gift from GENIE Project (Addgene plasmid # 234452; http://n2t.net/addgene:234452; RRID:Addgene_234452)). The vGlut1-pHluorin reporter was described previously **(Vevea & Chapman, 2023)**. We created a new GFAP-based FUGW plasmid replacing hSyn with GFAP for cytosolic mRuby3 expression and mouse APOE-HaloTag7 expression. We also made a hSyn-based cytosolic msGFP construct, and a synaptophysin-mRuby3 (SYP-mRuby3) lentiviral vector. All plasmids can be found deposited in Addgene.

### Lentivirus production and use

Lentiviruses were produced by the St. Jude Vector Laboratory. Lentiviral constructs were all based on the transfer plasmid FUGW. Hippocampal cultures were transduced with lentivirus at 1-5 DIV.

### Live cell imaging and quantification

Live cell fluorescence images were acquired on an Olympus IX83 inverted microscope equipped with an X-cite XYLIS LED (Excelitas; XT720S), Olympus 100x/1.5 NA objective (Evident; UPLAPO100XOHR), and an ORCA Fusion BT sCMOS camera (Hamamatsu Photonics) running Micro-manager software **(Edelstein, Amodaj, Hoover, Vale, & Stuurman, 2010)**. Standard imaging media (extracellular fluid; ECF) consisting of 140 mM NaCl, 3 mM KCl, 1.5 mM CaCl_2_, 1 mM MgCl_2_, 5.5 mM glucose, 10 mM HEPES (pH 7.3), B27 (Gibco), GlutaMAX (Gibco) were used for imaging experiments. For iGluSnFR and pHluorin stimulated release experiments, 50 μM D-AP5 (Tocris; 0106), 20 μM CNQX (Tocris; 1045), and 100 μM picrotoxin (Tocris; 1128) were added to the imaging media. For spontaneous iGluSnFR imaging, 1 μM tetrodotoxin (Tocris; 1069) was added to the imaging media. For iGluSnFR, single image planes were acquired with 10 ms exposure (100 Hz) using Sedat 490/20nm filter excitation and 525/36 nm filter emission. For pHluorin, single image planes were acquired with 500 ms exposure (2 Hz) using Sedat 490/20nm filter excitation and 525/36 nm filter emission. Single planes were selected so the highest amount of biosensor was in focus. For stimulated release imaging, stimuli were triggered by a Model 4100 Isolated High-Power Stimulator (A-M Systems; 930000) through platinum wires attached to a field stimulation chamber (Warner Instruments; RC-49MFSH), integrated by an Axon Digidata 1550B (Molecular Devices). Voltage for field stimulus was set to the lowest voltage that reliably produced presynaptic calcium transients in >95% presynaptic boutons using synaptophysin-GCaMP6f as a presynaptic calcium reporter. Experiments were performed at 34 °C. Temperature and humidity were controlled by an Okolab incubation controller and chamber. HTL-conjugated JF dyes were graciously provided by Luke Lavis from the Janelia Research Campus. Quantification of iGluSnFR4f **(Aggarwal et al., 2025)** and vGlut1-pHluorin was performed using a custom ImageJ macro described and used in **(Vevea & Chapman, 2020)** and **(Vevea & Chapman, 2023)**.

### Colocalization quantification

Pearson correlation coefficient (PCC) and Manders coefficients was quantified using Fiji for ImageJ **(Schindelin et al., 2012)** and Just Another Colocalization Plugin (JACoP) **(Bolte & Cordelieres, 2006)**, specifically, the BIOP JACoP implementation from the PTBIOP Fiji update site. APOE KO mouse brain tissue was used and prepared as described in ‘**Tissue Fixation, Sectioning, and Immunofluorescence**’ methods section. All images were captured using identical settings and the APOE KO images were used to estimate background intensity defined by measuring average intensity from APOE KO ROI plus 3x standard deviation. This value was subtracted from all WT APOE channels before costes automatic threshold and BIOP JACoP analysis.

### Chemicals

Chemicals are from Sigma-Aldrich unless otherwise specified and detailed in their methods section. All chemicals were resuspended and stored per manufacturer guidelines.

### Statistics

Statistics related to proteomics and lipidomics are described in their respective sections. All other analyses are detailed here. Exact values from experiments and analysis, including the number of data points (*n*) and trials (i.e., biological replicates) for each experiment, are listed in the figure legends. GraphPad Prism 11.0.0 (GraphPad Software) was used for statistical analysis. When appropriate, data are displayed as Tukey box and whisker plots. All statistical tests were two-sided unless otherwise noted. No *a priori* power analysis was completed before experimentation; however, to estimate sample size, a nomogram for sample size, effect size, and power was consulted **(Serdar, Cihan, Yucel, & Serdar, 2021)**. For our experimental planning, we generally prefer power = 0.8 and confidence = 0.05. Then, we estimate the standardized difference to arrive at a rough sample size result. For most experiments we chose a standardized difference = 1, representing a large difference expectation, and resulting in an estimated sample size of ∼30. For biosensor image analysis, many boutons were imaged. The *n* that was quantified was the average of all responding regions of interest (ROI) from a single field of view (FOV) and evenly dispersed between different image sets and biological replicates.

## Supplemental Figures and Legends

**Supplemental Figure 1:**
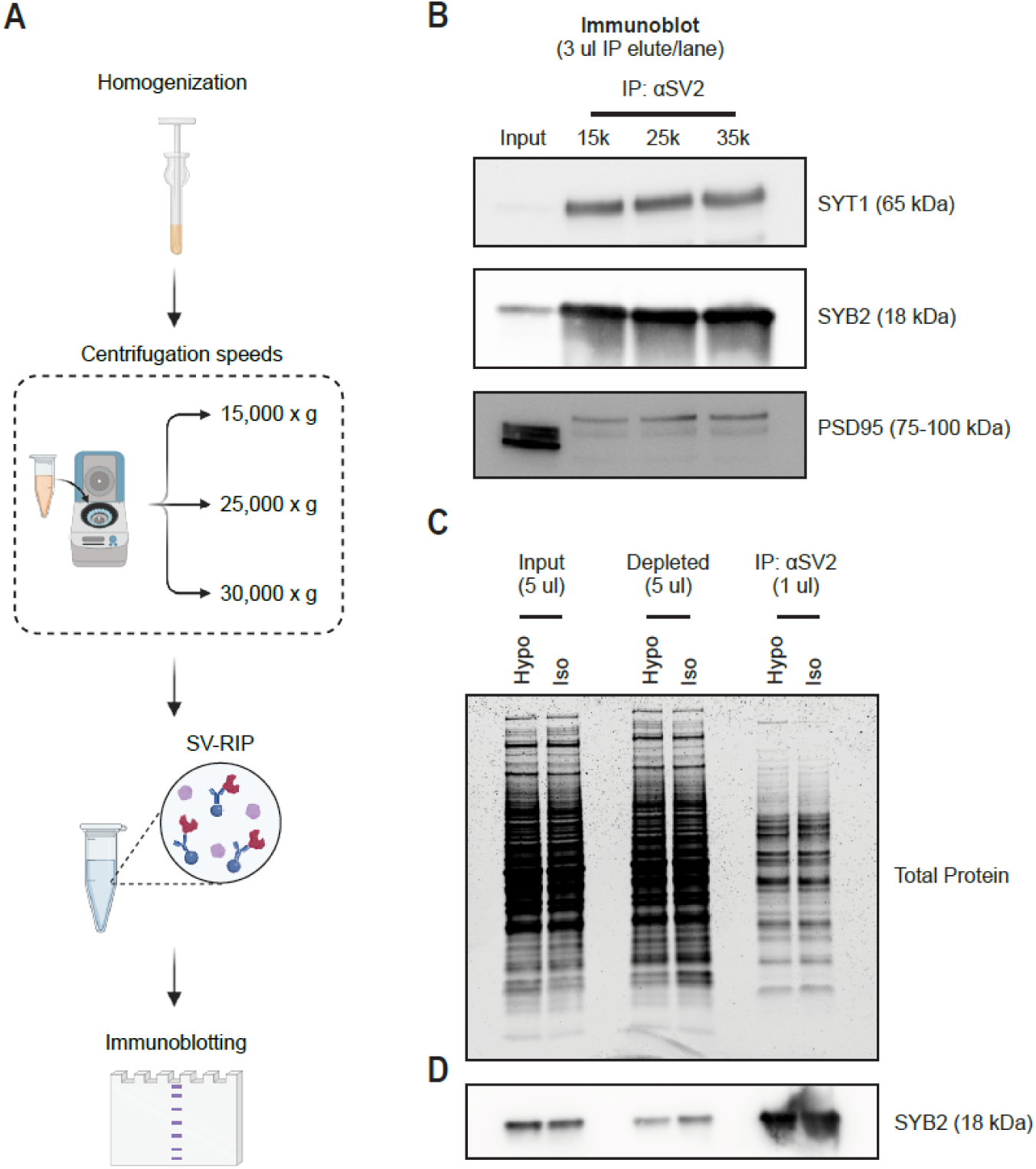
SV-RIP validation: Influence of centrifugation speeds on the purity of isolated synaptic vesicles. **A.** Experimental workflow for testing different centrifugation speeds during immunoprecipitation. After homogenization, the supernatant was centrifuged at 15,000, 25,000, or 30,000 x g and taken for SV-RIP. Samples were then immunoblotted to assess whether centrifugation speed alters protein composition. **B.** Immunoblot analysis of samples centrifuged at 15k, 25k, and 35k x g for synaptic vesicle markers (SYT1 and SYB2) and post-synaptic marker (PSD95) was performed to assess synaptic vesicle enrichment and purity with different centrifugation speeds. **C.** Total protein detected for Input, SV-depleted (Depleted), and SV-enriched (IP: αSV2) fractions using hypotonic or isotonic homogenization buffer. **D.** Immunoblot analysis of SYB2 for Input, Depleted, and IP: αSV2 fractions using hypotonic or isotonic homogenization buffer.

**Supplemental Figure 2:**
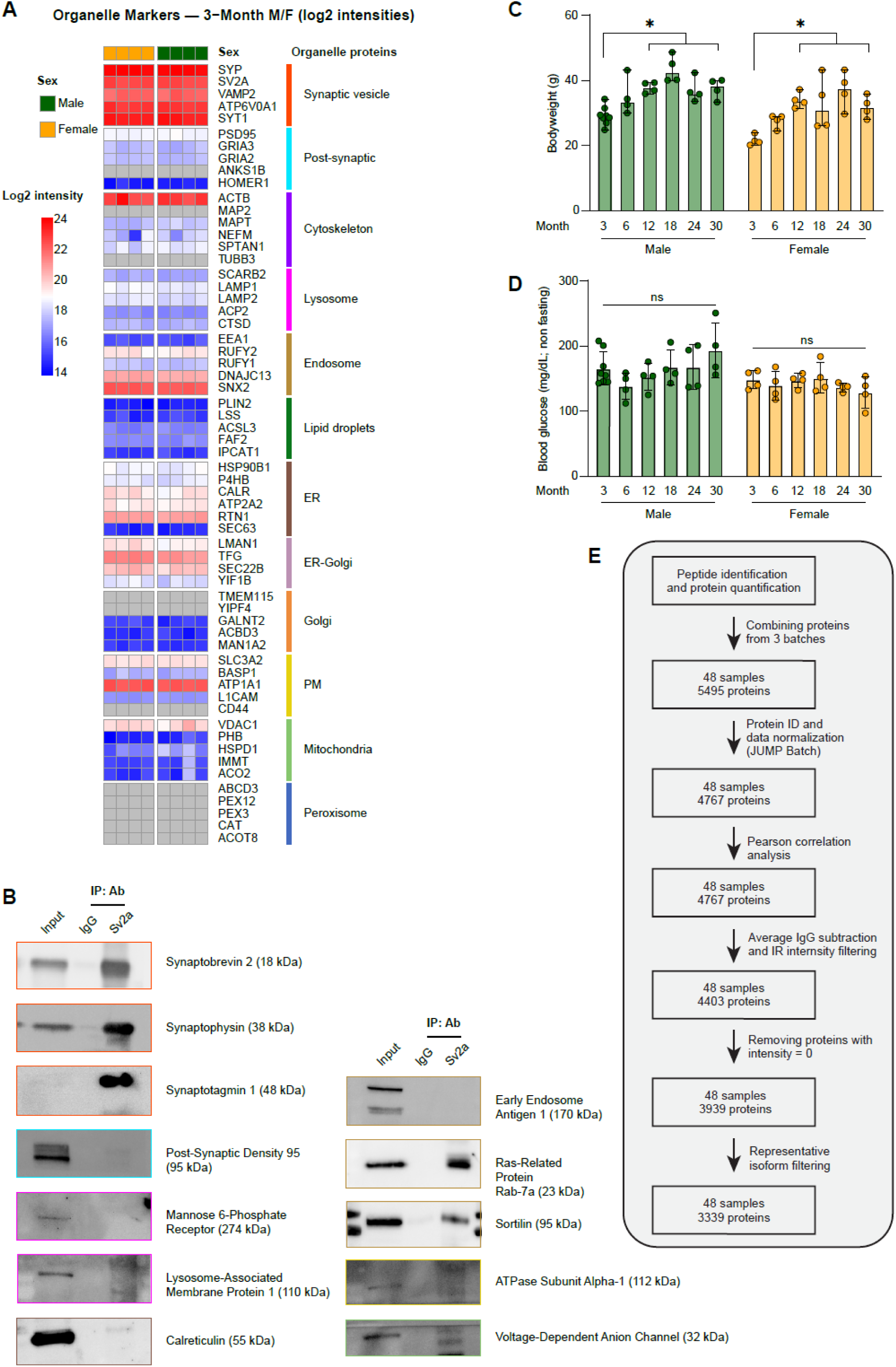
Additional validation of SV-RIP enrichment and mouse biomarker analysis across groups. **A.** Log2 intensity of organelle markers from 3-month male and female samples detected from TMT proteomics. **B.** Organelle markers examined by western blot from Input, IgG IP, and SV2a IP. **C.** Body weight (grams) at collection for male and female C57BL6/J mice. Data are displayed as median with 95% confidence intervals and analyzed by 2-way ANOVA with Dunnett’s multiple comparisons test (* p<0.05). **D.** Non-fasting blood glucose levels (mg/dL) at collection for male and female C57BL6/J mice. Data are displayed as median with 95% confidence intervals and analyzed by 2-way ANOVA with Dunnett’s multiple comparisons test. **E.** Proteomic data analysis workflow. Using the St. Jude JUMPm software, peptides were identified and quantified from 48 samples, and proteins detected in all three batches combined (5495 proteins). Protein IDs were assigned and data normalized using JUMP Batch (4767 proteins), and Pearson correlation analysis was performed. The average IgG intensity was subtracted for each protein, and those not detected in the Internal Reference (IR) samples removed (4404 proteins). Proteins with an intensity of 0 in any sample were removed (3939 proteins). Finally, proteins were filtered to retain one isoform, and 3339 proteins were taken for downstream statistical analysis.

**Supplemental Figure 3:**
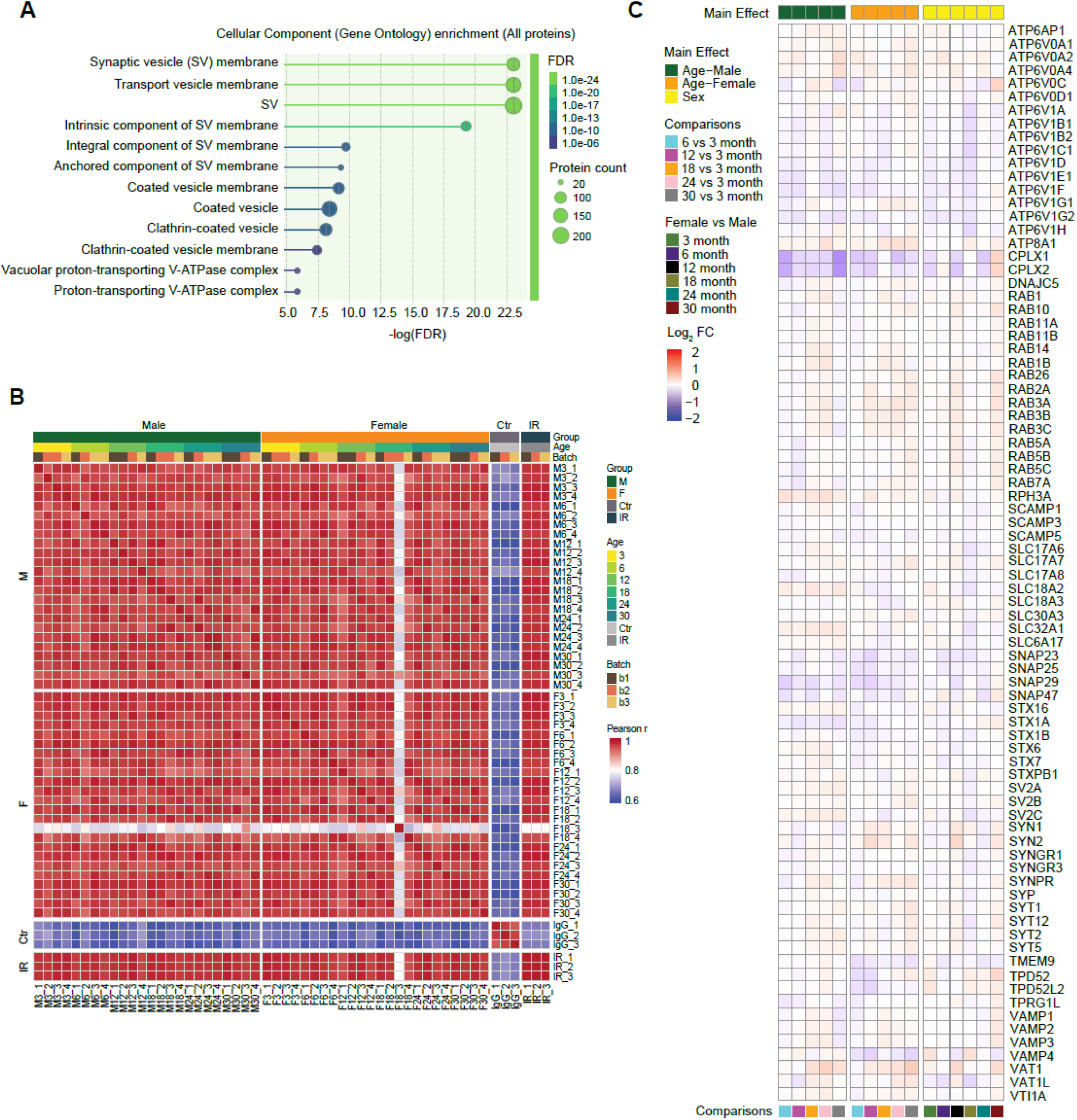
Additional proteomic data analysis. **A.** Cellular component gene ontology enrichment analysis of all detected proteins shows enrichment of synaptic vesicle components. **B.** Pearson correlation plot of 4764 normalized proteins from Male, Female, IgG Control (Ctr), and Internal Reference (IR) samples. **C.** Fold change of synaptic vesicle proteins for age in males (6 vs 3 month, 12 vs 3 month, 18 vs 3 month, 24 vs 3 month, and 30 vs 3 month), age in females (6 vs 3 month, 12 vs 3 month, 18 vs 3 month, 24 vs 3 month, and 30 vs 3 month), and sex (Female vs Male 3 month, 6 month, 12 month, 18 month, 24 month, and 30 month).

**Supplemental Figure 4:**
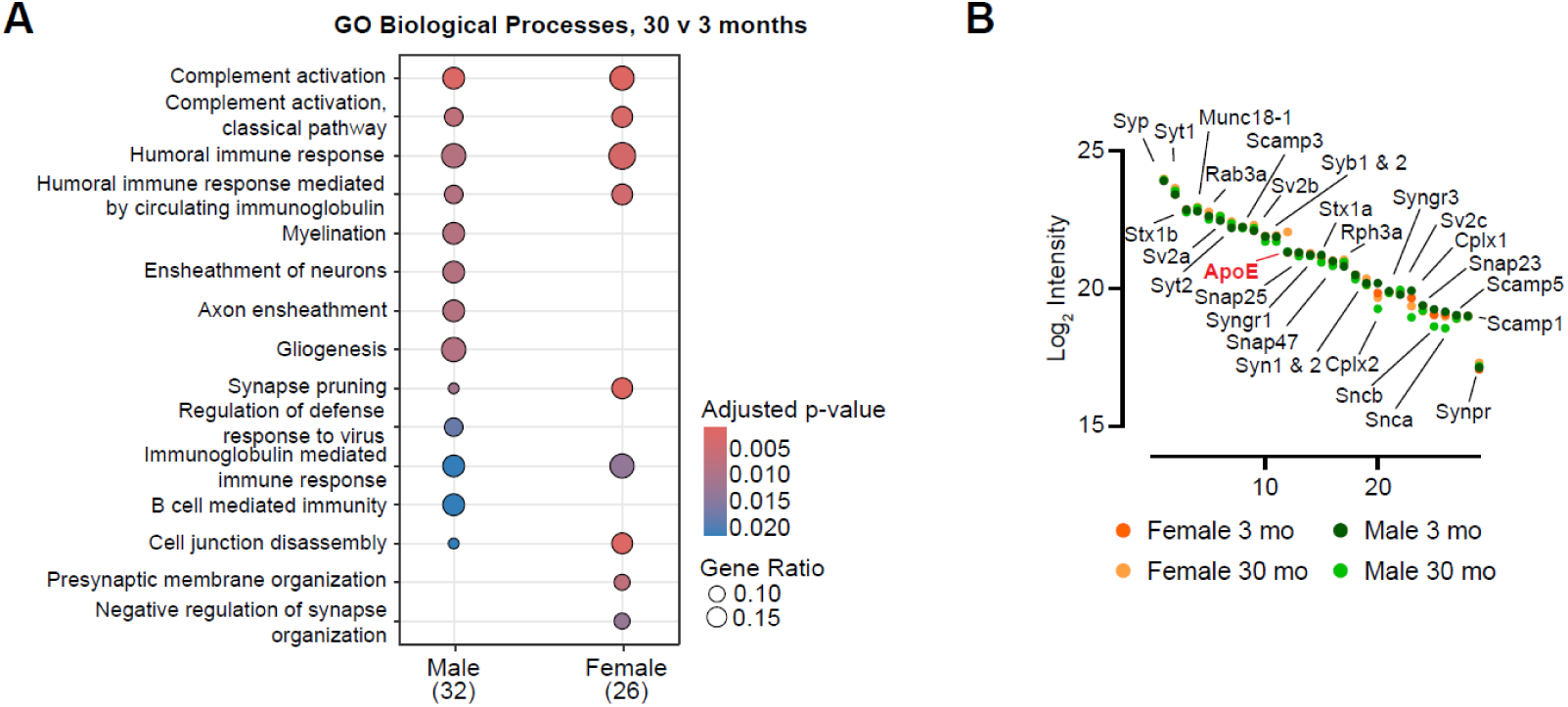
Additional analysis of proteomic changes between young and old SVs from male and females. **A.** Top GO Biological Processes for proteins significantly altered between 3 and 30 months for male and female samples. **B.** Rank ordered log2 intensity of Core SV proteins from 3- and 30-month-old males and females as shown in Main Figure 2E, now with levels of APOE added to chart demonstrating APOE is an abundant SV protein in males and females.

**Supplemental Figure 5:**
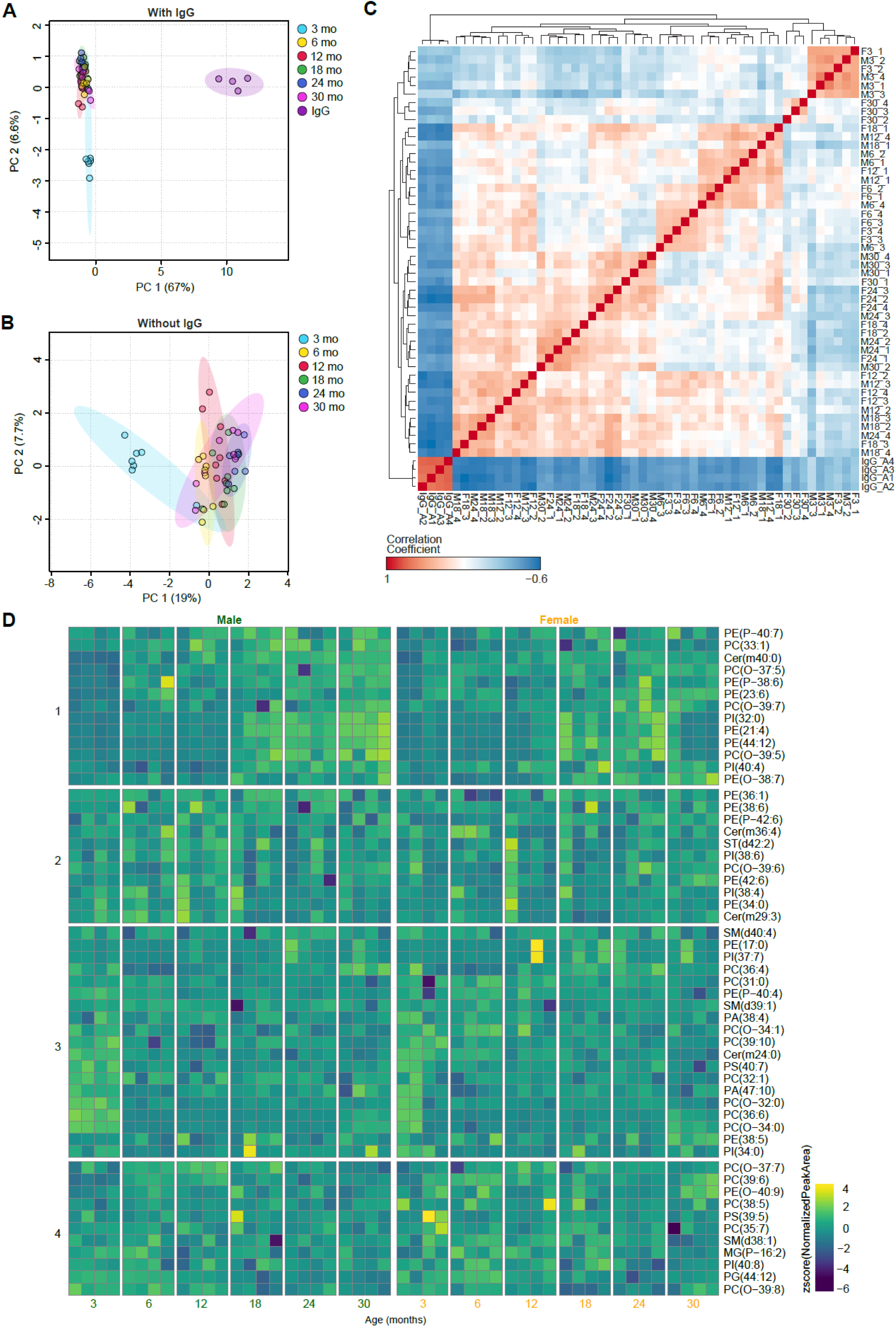
Lipidomic quality control and further analysis. **A.** Principal component analysis of detected phospholipids at all ages (sex combined) for IgG and SV samples. IgG samples separate from SV samples in the first component. **B.** Principal component analysis of detected phospholipids at all ages (sex combined) for SV samples only. 3 month samples separate from other ages in the first component. **C.** Correlation plot of phospholipids detected from untargeted lipidomics for all IgG and SV samples. IgG phospholipids are highly correlated, and distinct from SV phospholipids. **D.** Phospholipids with a significant Age and Sex interaction (FDR ≤0.05). Values displayed are z-score of normalized peak area for each significantly altered phospholipid from male and female synaptic vesicles at 3-, 6-, 12-, 18-, 24-, and 30-months. Data were analyzed by limma linear regression with Benjamini-Hochberg multiple testing correction and separated into 4 k-means clusters.

**Supplemental Figure 6:**
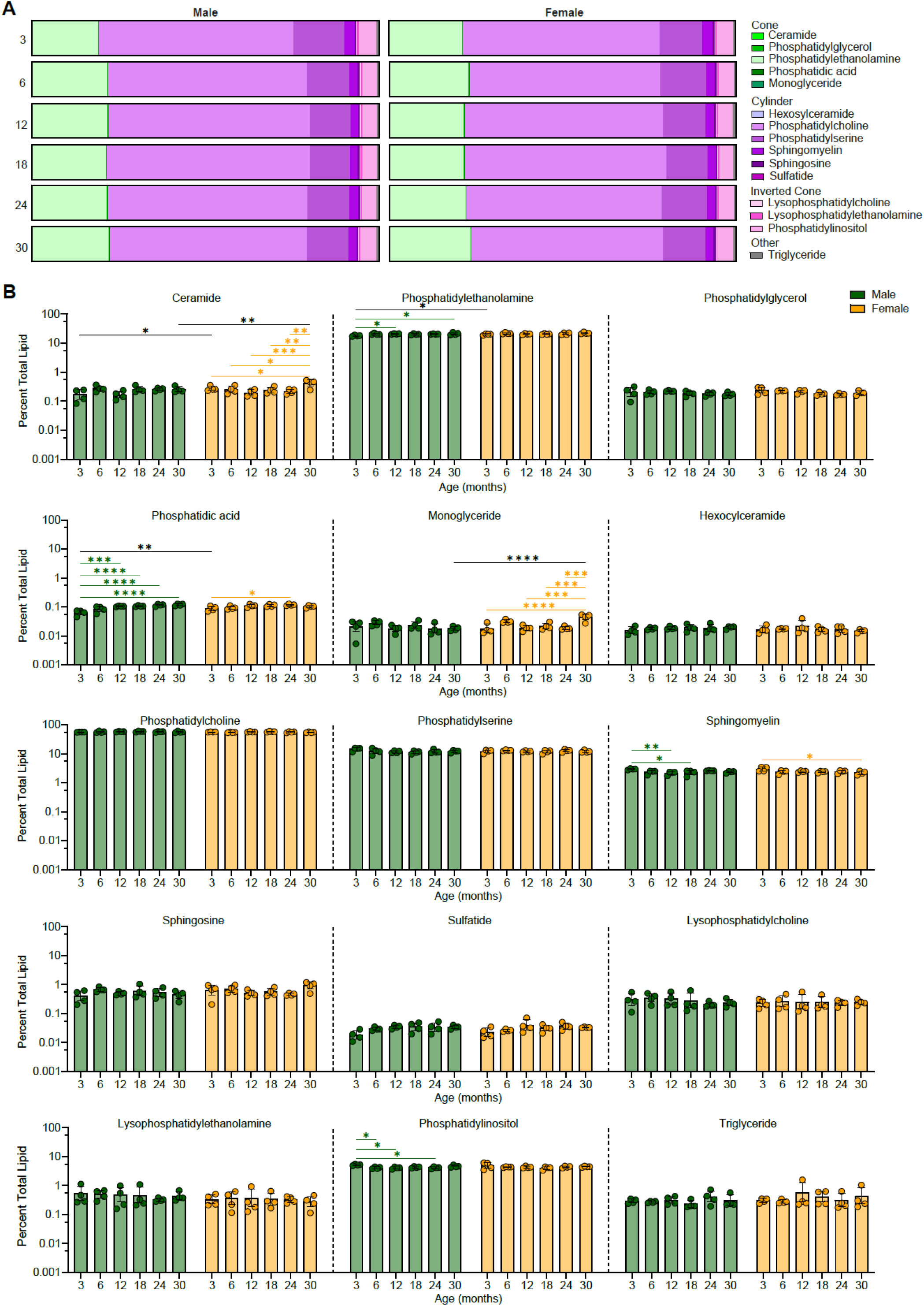
Phospholipid class analysis. **A.** Average percent total lipid composition by lipid class for male and female synaptic vesicles at 3-, 6-, 12-, 18-, 24-, and 30-months. **B.** Percent total composition of individual lipid classes (ceramide, phosphatidylethanolamine, phosphatidylglycerol, phosphatidic acid, monoglyceride, hexocylceramide, phosphatidylcholine, phosphatidylserine, sphingomyelin, sphingosine, sulfatide, lysophosphatidylcholine, lysophosphatidylethanolamine, phosphatidylinositol, and triglyceride) for male and female synaptic vesicles at 3-, 6-, 12-, 18-, 24-, and 30-months. Data are displayed as means with standard deviation and analyzed by 2-way ANOVA with Tukey’s multiple comparisons test (* p<0.05, ** p<0.01, *** p<0.001, **** p<0.0001).

**Supplemental Figure 7:**
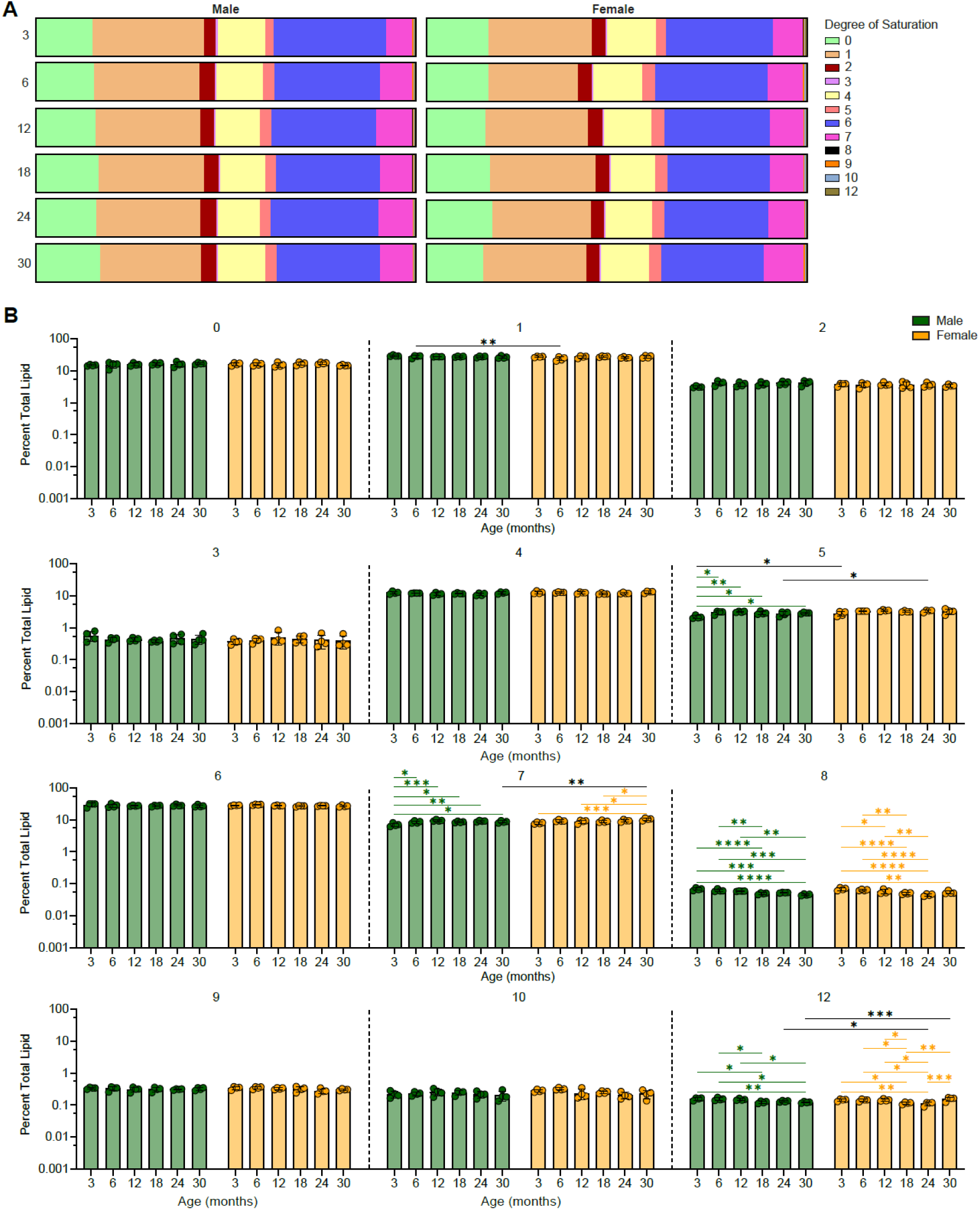
Phospholipid sidechain analysis. **A.** Average percent total lipid composition by lipid saturation for male and female synaptic vesicles at 3-, 6-, 12-, 18-, 24-, and 30-months. **B.** Percent total lipid composition by degree of saturation (0, 1, 2, 3, 4, 5, 6, 7, 8, 9, 10, 12 double bonds) for male and female synaptic vesicles at 3-, 6-, 12-, 18-, 24-, and 30-months. Data are displayed as means with standard deviation and analyzed by 2-way ANOVA with Tukey’s multiple comparisons test (* p<0.05, ** p<0.01, *** p<0.001, **** p<0.0001).

**Supplemental Figure 8:**
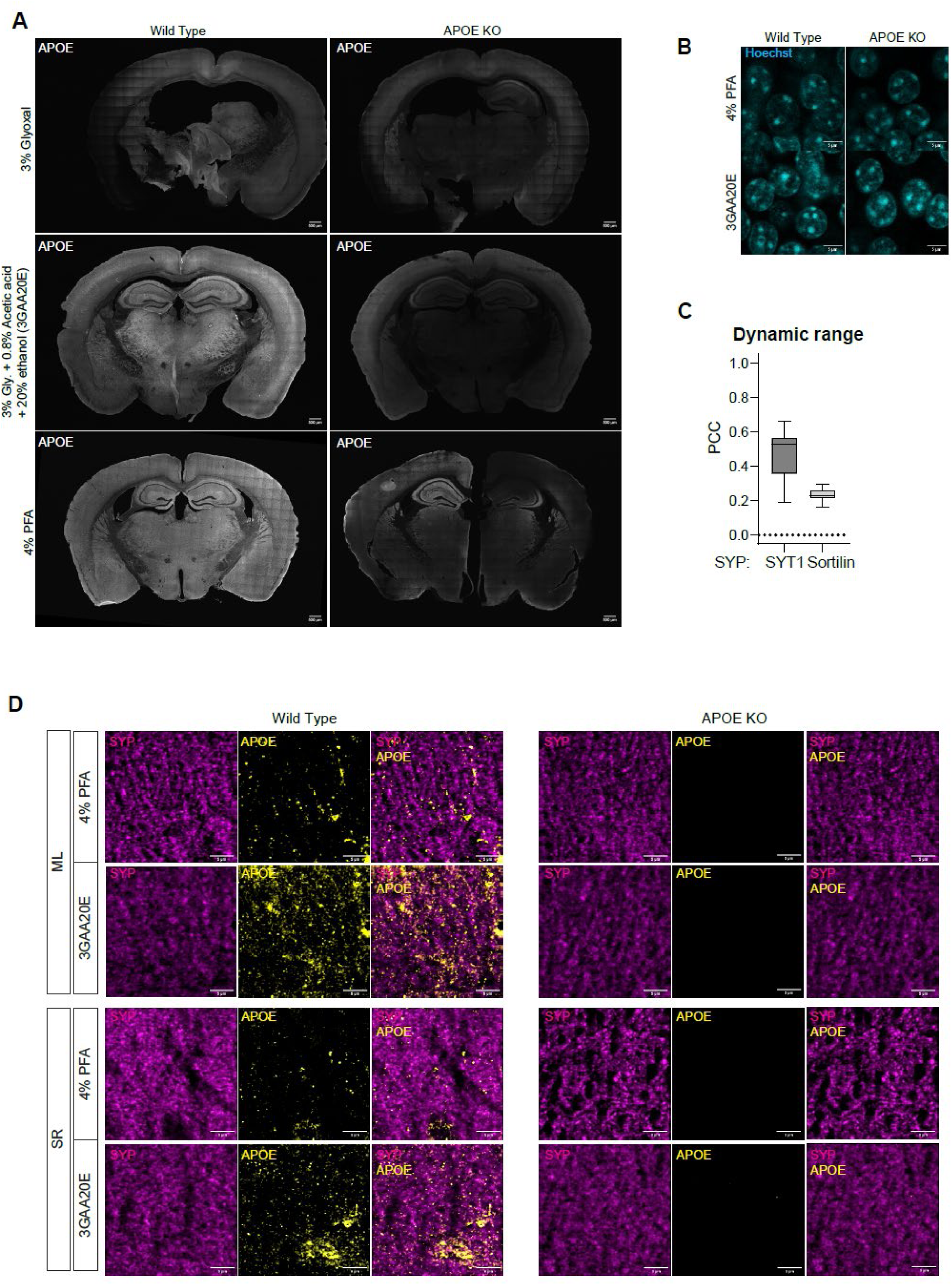
Brain tissue fixation optimization and comparison. **A.** Whole-section immunohistochemical comparison of APOE labeling in wild-type (left column) and APOE knockout (KO; right column) mouse brains fixed using three different fixatives including 3% glyoxal (top row), 3GAA20E (middle row), and 4% PFA (bottom row). **B.** Nuclear morphology and size appear comparable between two fixatives. Hoechst staining of wild-type (left column) and APOE KO (right column) mouse brains fixed with 4% PFA (top row) or 3GAA20E (bottom row). **C.** Dynamic range assessment of the colocalization analysis using Pearson’s correlation coefficient (PCC). Positive (SYP-SYT1) and negative (SYP-Sortilin) marker pairs were stained to demonstrate that the analysis method reliably distinguishes true synaptic colocalization from non-colocalizing signals. **D.** Comparison of APOE immunostaining quality in the molecular layer (ML) and stratum radiatum (SR) of APOE wild-type and knockout mouse hippocampi fixed with 4% PFA (top row) or 3GAA20E (bottom row). Anti-APOE labeling (yellow) in the KO tissue serves as a negative control, revealing reduced background fluorescence and improved signal-to-noise with 3GAA20E fixation relative to PFA. The anti-SYP staining (magenta) highlights synaptic structures present in both conditions.

**Supplemental Figure 9:**
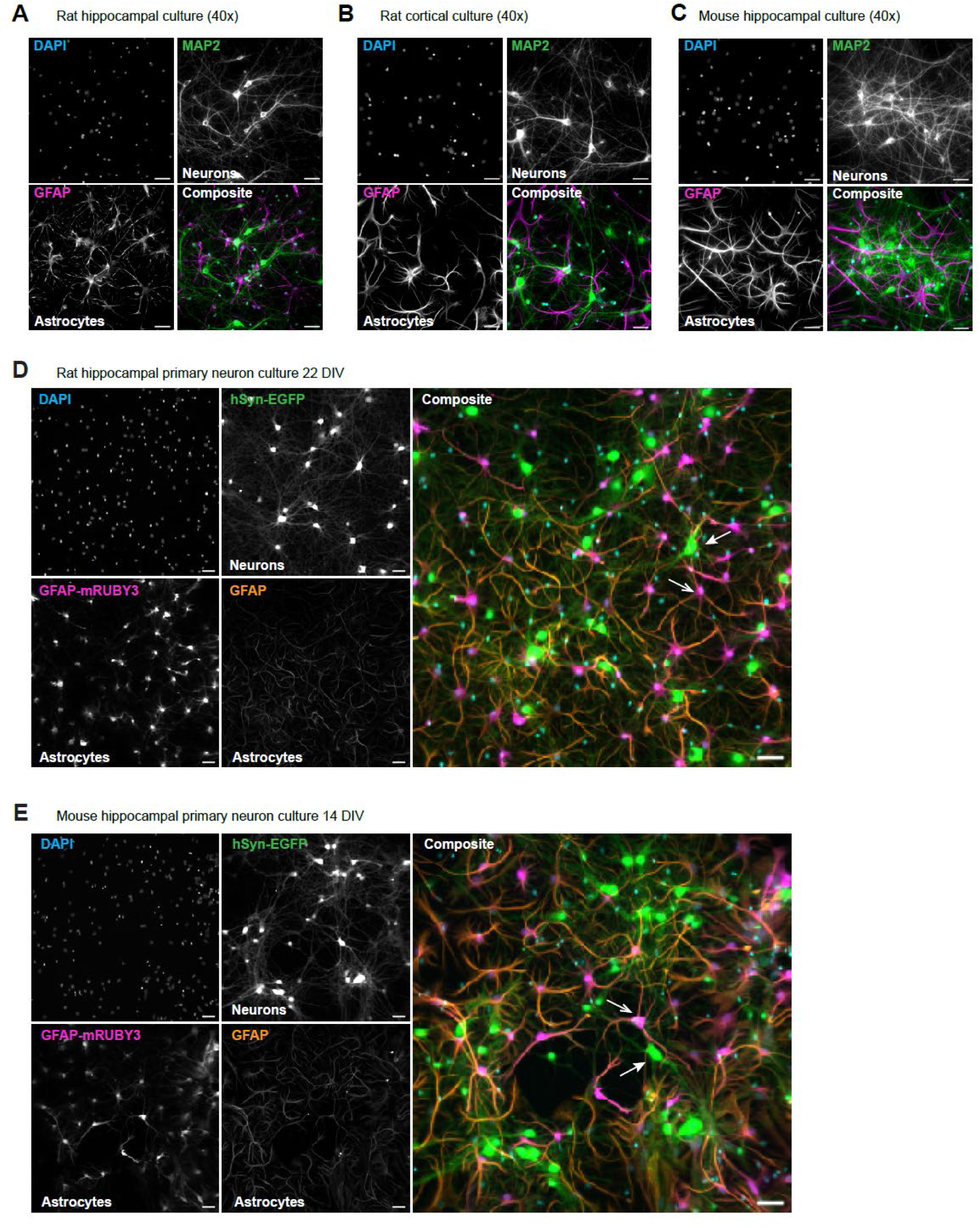
Validation of mixed neuron-astrocyte co-cultures. **A.** Representative immunofluorescence image demonstrating the presence of both neurons and astrocytes in DIV21 rat hippocampal cultures. Neurons are visualized with MAP2 (green), astrocytes with GFAP (magenta) and nuclei with DAPI (blue). **B.** Representative immunofluorescence visualization confirming the presence of neurons and astrocytes in DIV21 rat cortical cultures. Neurons are marked by MAP2 (green), astrocytes by GFAP (magenta), and nuclei by DAPI (blue). **C.** Representative immunofluorescence image illustrating neuronal and astrocytic populations in DIV14 mouse hippocampal culture labeled with MAP2 and GFAP. Stained with anti-MAP2 (green), anti-GFAP (magenta), and DAPI (blue). **D.** Lentiviral delivery of a *GFAP: cytosolic mRUBY3* construct at high titer drives reporter expression selectively in astrocytes in DIV22 rat hippocampal culture. Neurons exhibit robust hSyn-EGFP expression (green), while astrocytes display strong GFAP: cytosolic mRUBY3 fluorescence (orange). **E.** In DIV14 mouse hippocampal cultures, high titer of hSyn: cytosolic msGFP and GFAP: cytosolic mRUBY3 lentiviruses drive cell-type-specific labeling, with mRUBY3 restricted to GFAP-positive astrocytes and absent from neurons, validating promoter specificity.

**Supplemental Figure 10:**
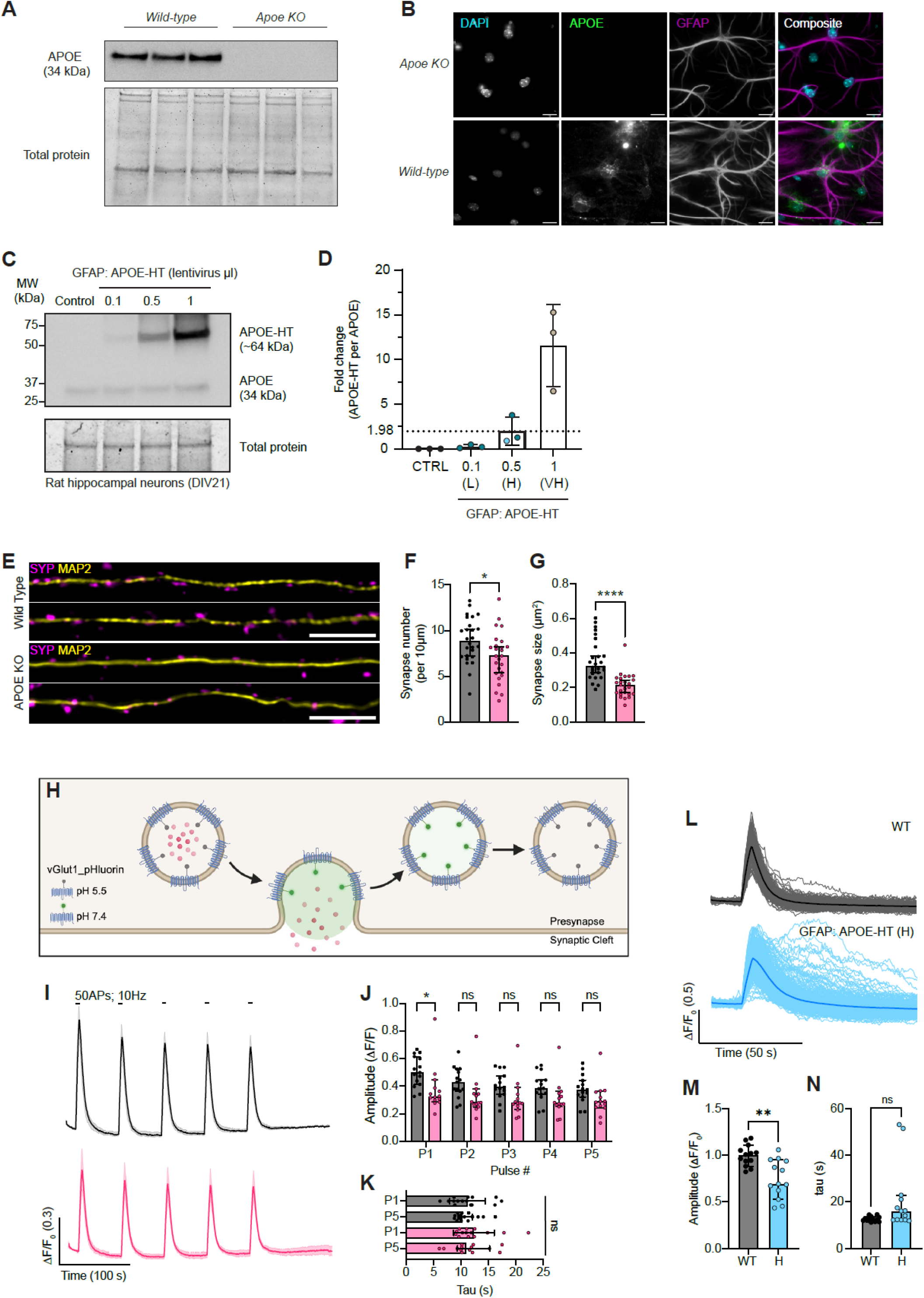
Modulation of APOE protein levels and synaptic characterization. **A.** Representative immunoblot from 3 separate DIV14 hippocampal cultures from either wild type or APOE KO mice probed for APOE. **B.** Representative immunofluorescence image localizing APOE in wild-type and APOE KO DIV14 mouse hippocampal neuron astrocyte cocultures. Stained with DAPI (blue), anti APOE (green), and anti GFAP (magenta). **C.** Representative immunoblot from WT rat hippocampal DIV21 neuron astrocyte cocultures from control, and GFAP: APOE-HaloTag transduced cultures using 0.1 µl lentivirus (Low; L), 0.5 µl lentivirus (High; H), or 1.0 µl lentivirus (Very High; VH). **D.** Quantification of triplicate repeat immunoblots as shown in (F). **E.** Primary mouse hippocampal cocultures fixed and stained with anti MAP2 (yellow) and anti SYP (magenta). MAP2 regions straightened using ImageJ. Scale bar 5 microns. **F.** Synapse number (SYP punctae) per 10 microns. Values are median ± 95% confidence interval, n = 26 dendrite stretches per condition, Mann-Whitney two-tailed test, p < 0.05. **G.** Synapse size (SYP area) between WT and APOE KO cocultures. Values are median ± 95% confidence interval, n = 26 dendrite stretches per condition, Mann-Whitney two-tailed test, p < 0.0001. **H.** Schematic of vGlut1-pHluorin during exocytosis and endocytosis. **I.** Average trace of vGlut1-pHluorin with 50 action potentials; 10Hz train stimulus repeated 5 times with 1 minute rest in between trains; traces are mean± 95% confidence interval. **J.** Amplitude of evoked pHluorin events (ΔF/F_0_) from WT and APOE KO hippocampal cultures. Values are median ± 95% confidence interval, n = 14-15 neurons per condition. Two-way ANOVA with Sidak’s multiple comparisons test (* p<0.05) **K.** Time constant (tau) of the decay of first and fifth pHluorin peaks for WT and APOE KO hippocampal cultures. Values are median ± 95% confidence interval, n = 14-15 neurons per condition. One-way ANOVA with Dunnett’s multiple comparisons test. **L.** Representative traces from a single field of view (FOV) of Rat hippocampal neuron astrocyte cocultures expressing vGlut1-pHluorin and stimulated with a high frequency train of 100 action potentials at 20 Hz. WT in black and GFAP: APOE-HT OE (0.5 µl, High) in blue/light blue. Individual traces in light colors with FOV average in dark. **M.** Peak amplitude (ΔF/F_0_) of vGlut1-pHluorin from WT and GFAP: APOE-HT OE (0.5 µl, High). Values are median ± 95% confidence interval, n = 13 FOV per condition, Mann-Whitney two-tailed test, p < 0.005. **N.** Time constant (tau) of vGlut1-pHluorin decay after stimulation for WT and GFAP: APOE-HT OE (0.5 µl, High). Values are median ± 95% confidence interval, n = 13 FOV per condition, Mann-Whitney two-tailed test, ns = 0.0501.

